# Interpreting computational mechanism from multi-area neural recordings

**DOI:** 10.64898/2026.11.09.765426

**Authors:** Christopher Versteeg, Jonathan A. Michaels, Mehrdad Kashefi, Ayesha Vermani, Trung Le, J. Andrew Pruszynski, Chethan Pandarinath

## Abstract

Distilling computational mechanisms from neural recordings requires disambiguating intrinsic dynamics from external inputs. Here, we introduce Dynamically Interacting Systems (DynISys), a multi-compartment dynamical-systems model that infers per-area dynamics and structured inter-area communication from multi-area spiking activity. DynISys combines neural ordinary differential equation (ODE) latent dynamics with injective latent-to-neural mappings to achieve expressive and trustworthy dynamical inference. When applied to simultaneous dorsal premotor (PMd) and primary motor (M1) recordings from a continuous reaching task, DynISys accurately reconstructs neuronal responses without inventing spurious latent features. Analyzing the model’s dynamics reveals putative mechanisms for motor planning and execution, with PMd-to-M1 inputs positioning a stable fixed point in M1 that encodes execution dynamics for the upcoming movement, subsequently evoked by external (non-PMd) inputs. Finally, multi-session DynISys achieves consistent latents when fit to neural recordings spanning multiple days. In summary, DynISys allows distillation of multi-area recordings into an interpretable account of distributed neural computation.

## Introduction

A central goal of systems neuroscience is to understand how neural circuits drive behavior through their produced activity patterns (Shenoy et al. 2013; Vyas et al. 2020). Under the computation-through-dynamics view, a neural circuit’s dynamics (i.e., the rules that govern how its activity evolves over time) are a key substrate for computation (Churchland et al. 2012). Recent work has extended this view, highlighting that computation often emerges from complex interactions between local network dynamics and external inputs that modulate those dynamics (Sauerbrei et al. 2020, Inagaki et al. 2022, Driscoll et al. 2024, Oby et al. 2025). However, recordings of neural activity reveal neither a circuit’s intrinsic dynamics nor its external inputs directly. Generative models have sought to infer latent dynamics from these recordings (Pandarinath et al. 2018; Kim et al. 2021; Durstewitz et al. 2023; Linderman et al. 2016; Macke et al. 2011; Gao et al. 2016; Mastrogiuseppe and Ostojic 2018; Karniol-Tambour et al. 2024), with some also simultaneously inferring external inputs (Pandarinath et al. 2018; Schimel et al. 2022; Vahidi et al. 2025). However, as the same observed activity can be generated by many different combinations of dynamics and inputs, models may attribute intrinsic dynamics to fictitious inputs (or vice versa) (Versteeg et al. 2026, Fig. 1A,B).

**Figure 1.**
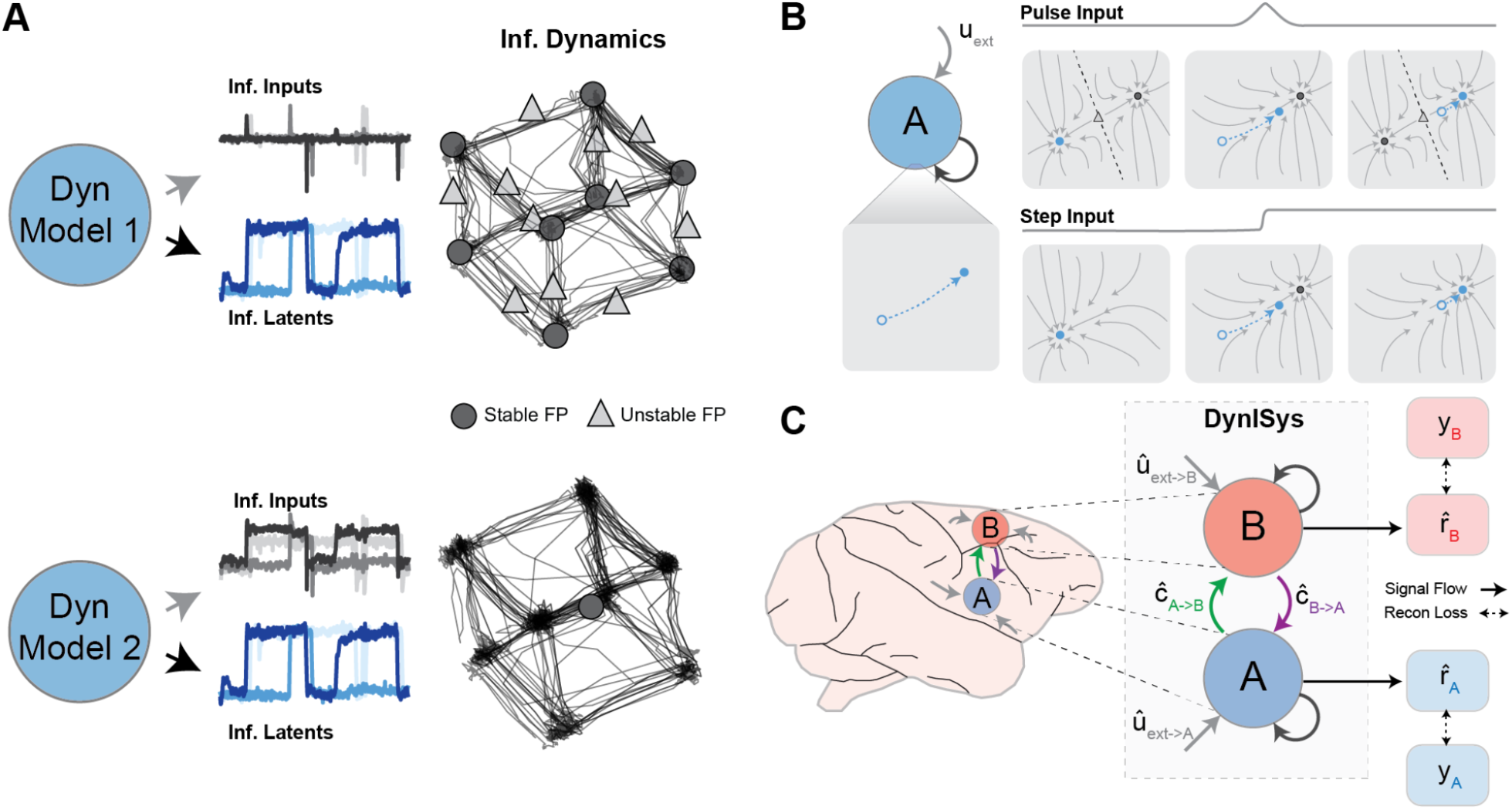
Disambiguating intrinsic dynamics and external inputs with interacting dynamics models. **(A)** Two example dynamics models from CtDToolkit (Versteeg et al. 2026) fit to synthetic 3-bit flip flop data, using different priors to constrain inferred external inputs (top: sparse prior; bottom: smooth autoregressive prior). Despite inferring similar latent activity (blue traces), the models infer different inputs (grey traces) and markedly different dynamical structures (latent trajectories and fixed points shown at right). Circles and triangles denote stable and unstable fixed points, respectively. **(B)** Schematic illustrating how different combinations of dynamics and inputs can generate the same latent trajectory. In one case (top), a transient pulse briefly reshapes the flow field, moving the state between two stable fixed points before the original flow field returns. In another case (bottom), a sustained step input instead shifts the stable fixed point yet produces an identical latent trajectory. Grey arrows indicate flow fields; blue markers and dotted arrows indicate the evolving latent state. **(C)** Left: Schematic of two brain areas A and B. Each area has intrinsic dynamics, inter-area communication (green, purple) and inputs from other brain areas (grey). Right: DynISys models each brain area as a separate latent system that evolves over time according to its own intrinsic dynamics, communication from other recorded brain areas (denoted by ĉ), external inputs (denoted by û), and an injective function that maps inferred latent activity onto predicted firing rates. DynISys is trained to predict firing rates 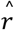 that optimally reconstruct recorded multi-area neural activity *y* as observations of a low-D interacting dynamical system. Subscripts indicate the relevant brain area for each variable: e.g., ĉ_A_ indicates all inferred communication signals to area A. Solid lines indicate a signal flow, while dashed lines indicate the reconstruction training objective.

Simultaneous recordings from multiple brain areas could provide more reliable accounts of neural computation by helping disambiguate the relative contributions of inputs and intrinsic dynamics. However, previous methods for inferring dynamics and inter-area communication jointly from simultaneously recorded populations (Glaser et al. 2020; Perich et al. 2026; Liu et al. 2025) typically have excess dimensions (Sedler et al. 2023) and are prone to hallucinations that reflect modeling artifacts rather than features of the underlying circuit (Sedler et al. 2023, Versteeg et al. 2024). For an interacting systems model to be interpretable (amenable to mechanistic interrogation), it must be both *expressive* (able to capture the rich dynamics of each area) and *trustworthy* (resistant to spurious dynamical features).

We build on previously published dynamical inference models to develop Dynamically Interacting Systems (*DynISys*; Fig. 1C). DynISys achieves expressive, low-dimensional latent dynamics via neural ordinary differential equation (ODE) networks (Chen et al. 2018; Kim et al. 2021; Sedler et al. 2023, Versteeg et al. 2024), and minimizes hallucinations in the modeled dynamics via injective readouts (Versteeg et al. 2024). Critically, DynISys has multiple compartments to constrain the relative contributions of intrinsic dynamics and inputs: each recorded area carries its own dynamics and the inputs exchanged between them are inferred explicitly. We apply DynISys to simultaneously recorded dorsal premotor (PMd) and primary motor (M1) cortical populations from two monkeys performing a sequential reaching task (Michaels et al. 2026 (in prep.)). We show that DynISys reproduces the response properties of recorded neurons while being markedly more resistant to invented features than alternative multi-compartment models. The timing and directionality of the inferred communication recapitulate the appearance of target information in PMd and M1 recordings and are consistent with a motor plan being transmitted from PMd to M1 during motor preparation.

As an *in-silico* model of circuit dynamics, DynISys allows researchers to test mechanistic hypotheses that are difficult to evaluate from recordings alone. To provide a proof-of-principle for this approach, we ask how inputs to M1 select and initiate motor plans. Analysis of the inferred M1 flow field during motor preparation reveals a family of input-dependent fixed points whose locations encode features of the upcoming reach (Inagaki et al. 2019; Shah et al. 2025). Targeted *in-silico* perturbations reveal that PMd inputs determine the preparatory fixed-point location on each trial and that these fixed points serve as initial conditions for subsequent execution dynamics triggered by external inputs to M1. Finally, we show that DynISys can aggregate data from multiple experimental sessions into a single dynamics model with consistent latent activity despite a changing set of recorded neurons. Together, these results illustrate how DynISys can translate multi-area recordings into candidate computational mechanisms and experimentally testable predictions.

## Results

### DynISys captures behaviorally salient features of PMd/M1 activity during a sequential reaching task

Our goal is to use dynamics models to uncover how interacting brain areas give rise to functional computation. We chose to analyze a sequential reaching task in which dynamics related to planning and execution must be coordinated by interacting neural populations. Monkeys performed a continuous reaching task in which they moved a cursor to a series of visual targets using a Kinarm manipulandum (Scott 1999) while we recorded neural activity simultaneously from PMd and M1 using Neuropixels probes (Fig. 2A, Monkey P: 64 and 63 single units, respectively; Monkey M: 28 and 13 single units, respectively; Michaels et al. 2025, Kashefi et al. 2026). Critically, the task includes trials in which the monkey sees both the current and upcoming targets (Fig. 2B), allowing them to plan the next reach while completing the current one. This planning during ongoing movement can help dissociate neural activity associated with preparation and execution.

**Figure 2.**
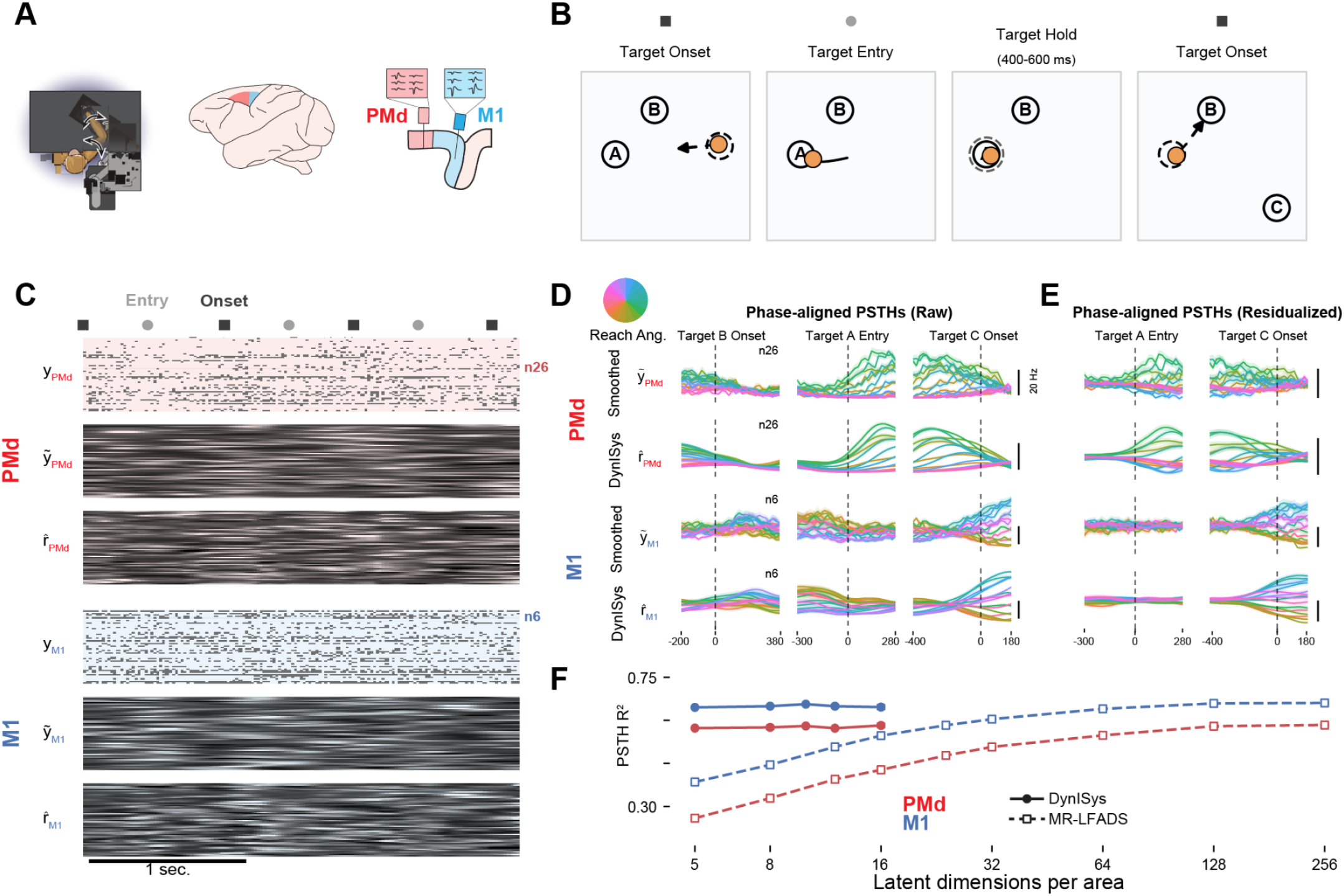
DynISys captures PMd/M1 population activity patterns during a sequential reaching task. **(A)** Kinarm behavioral apparatus and PMd/M1 recording-array locations. **(B)** Target (unfilled circles) and hand position (orange circle) across task phases of a single trial: target onset (target B appears, reach to target A begins), target entry (hand enters A), target hold, and second target onset (target C appears, reach to B begins). **(C)** Example PMd/M1 spike rasters (top), smoothed neural activity (middle; symmetric Gaussian kernel, σ = 100 ms) and matching DynISys-predicted firing-rate heatmaps (bottom) for the same stitched trial window, showing single-trial, per-channel rate agreement with observed spiking. **(D)** PMd/M1 PSTHs aligned to target B onset, target A entry, and target C onset, comparing observed and DynISys-predicted firing rates across phases. Reach angle indicated by colorwheel. **(E)** Same as (D), with PSTHs computed after removing components of the neural activity that were predictive of the current target location using linear models. **(F)** Reach direction PSTH R^2^ for DynISys and MR-LFADS as a function of latent dimensionality.

We trained DynISys to model the two recorded populations as a coupled dynamical system, with one compartment per area (as in Fig. 1). Each compartment carries its own latent state 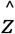 that evolves under a learned neural-ODE generator 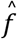, and the two compartments exchange information only through explicit, low-dimensional communication channels ĉ. Separate sets of inferred external inputs û account for drive that originates outside the recorded brain areas. Each compartment’s latent state is mapped onto high-dimensional firing rate predictions 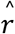 for that compartment via a nonlinear, injective normalizing-flow readout ĝ (see Methods for full implementation).

We trained models across latent dimensionalities and hyperparameter settings using a regularized Poisson reconstruction objective and selected a representative model based on reconstruction performance (Methods). The main findings were consistent across a range of hyperparameter settings. After restitching predictions into continuous recordings, the inferred rates tracked fluctuations in PMd and M1 activity and aligned with behavioral events, despite training without behavioral supervision (Fig. 2C).

We first assessed whether DynISys’s low-dimensional representation could reproduce target-related response patterns in recorded neurons. In this task, targets were displayed one reach in advance: the appearance of B cued the reach to A, and the appearance of C cued the reach to B. Between these cues, the monkey reached A and held there, with A as the current target and B visible as the upcoming target. To track B’s encoding over time, we grouped trials by the A-to-B reach angle and computed PSTHs aligned to B onset, entry into A, and C onset.

DynISys reproduced the resulting direction-dependent modulation in example PMd and M1 neurons (Fig. 2D). Across neurons, mean PSTH R^2^ was 0.61 in PMd and 0.66 in M1 for Monkey P (Monkey M: Extended Data Fig. 1). To reduce modulation associated with target A’s position, we removed the component linearly explained by its absolute coordinates from both observed and inferred activity before recomputing the PSTHs (Fig. 2E; Methods). In the position-adjusted examples, direction-dependent modulation emerged earlier in PMd than in M1, motivating the population-level timing analyses below. DynISys also captured responses associated with other behavioral features such as starting location and peak hand speed (Extended Data Figs. 2 and 3).

A key advantage of DynISys is that its dynamical expressivity is not tied to the dimensionality of its latent state. To assess the practical benefit of this flexibility, we compared PSTH R^2^ for DynISys against MR-LFADS models trained across a range of latent dimensionalities (Fig. 2F, see Methods, ‘Baseline models’, Liu et al. 2025). MR-LFADS reconstruction accuracy improved with increasing dimensionality, whereas DynISys achieved near peak PSTH R^2^ with only 8 latent dimensions per area, matching the performance of MR-LFADS models with 128 dimensions per area (0.60, 0.67 for 8-D DynISys PMd, M1 versus 0.59, 0.68 for 128-D MR-LFADS PMd, M1, respectively). We also evaluated an alternative multi-area dynamics model, Current-Based Decomposition (CURBD, Perich et al. 2026), but were unable to obtain model fits that were substantially better than predicting the mean firing rate of each neuron (see Methods, ‘Baseline models’). Crucially, DynISys captures diverse neural responses while retaining a compact latent space suited for dynamical analysis.

### DynISys infers trustworthy latent state estimates and communication signals consistent with transmission of a motor plan from PMd to M1

While DynISys captures complex structure from neural recordings, this does not guarantee that inferred latent features or dynamics are trustworthy, as flexible models often hallucinate latent structure that does not actually exist in the underlying circuit (Versteeg et al. 2026). To evaluate whether such hallucinations exist in DynISys, we tested whether the latents contained information about upcoming targets before that information was detectable in the observed spiking activity.

We first characterized the time course of target encoding in the neural recordings themselves by quantifying the decoding accuracy of the upcoming target over time (Fig. 3A). Decoding from PMd neural activity rose above chance well prior to current target entry, while decoding from M1 neural activity did not become significant until around the time of current target entry. We performed the same decoding analysis on the DynISys-inferred latent activity and found a strong temporal correspondence (Fig. 3A, Monkey M: Extended Data Fig. 4). This lends credibility to the inferred latents as trustworthy accounts of the state of the underlying neural circuit, a key prerequisite for any interrogation of these models for computational mechanisms.

**Figure 3.**
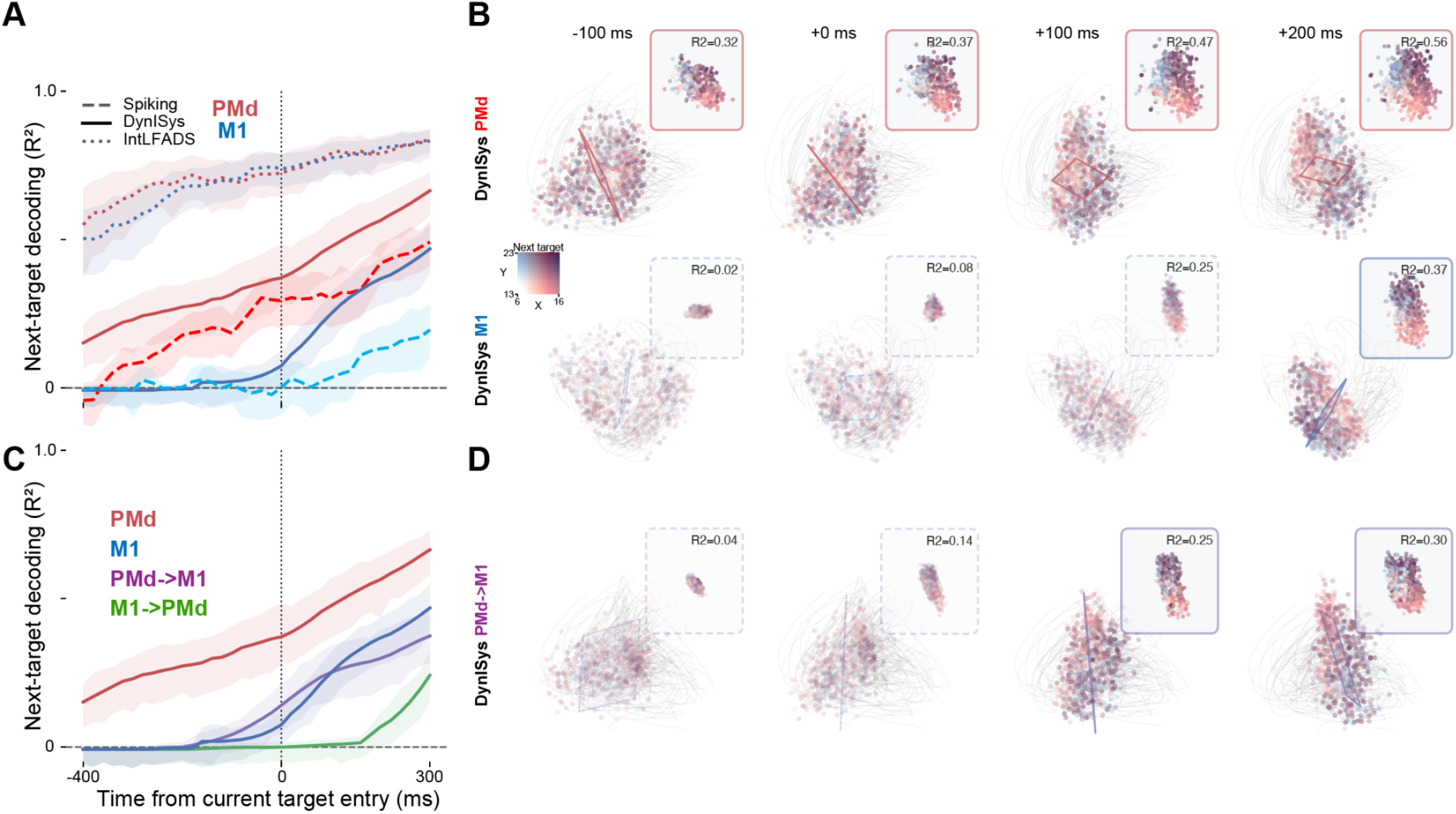
DynISys infers time course of target encoding within each area and transmission of target information across areas. **(A)** Time-resolved next-target decoding from PMd latents, PMd smoothed spikes, M1 latents, M1 smoothed spikes, and IntLFADS controls, aligned to current-target entry. Shaded areas indicate bootstrapped 95% confidence intervals for mean decoding performance. **(B)** State-space snapshots of PMd and M1 DynISys latents (top-3 PCs) at four times relative to current-target entry, each with a fitted linear decoding plane for next-target position (colored by target, R^2^ noted; solid border indicates performance significantly above chance); inset shows decoded-vs-true target projection per time slice. **(C)** Time-resolved next-target decoding from PMd→M1 communication vs. the M1→PMd control direction (plus PMd/M1 latents for reference). **(D)** Same state-space/plane treatment as (B) for the PMd→M1 communication channel.

We compared DynISys’ trustworthiness to two alternative models: an interacting-dynamics LFADS developed as a control for this study (IntLFADS) and a previously published multi-compartment dynamics model (MR-LFADS; Liu et al. 2025). We summarize the major differences in the candidate architectures in Extended Data Table 1.

**Table 1:** Kinarm dataset and preprocessing parameters.

| **Parameter/Monkey** | **Monkey P** | **Monkey M** |
|---|---|---|
| Spike bin width | 20 ms | 20 ms |
| Modeled trial window | 80 bins (1600 ms) | 80 bins (1600 ms) |
| Minimum mean firing rate | 1 Hz | 1 Hz |
| Held-in / held-out split | 80 % / 20 % | 80 % / 20 % |
| M1 single units (total / held-in) | 63 / 51 | 13 / 11 |
| PMd single units (total / held-in) | 64 / 52 | 28 / 23 |

The IntLFADS control extends latent factor analysis via dynamical systems (LFADS; Pandarinath et al. 2018) to multiple interacting neural populations without introducing any architectural constraints designed to ensure trustworthiness. Applying the same decoding analysis, we found that target information became decodable from IntLFADS latents well before it could be detected in recorded neural activity. In contrast with IntLFADS, MR-LFADS infers initial conditions causally and obligates inferred inter-area communication to affect predicted neural activity. However, its linear readout does not obligate that *all* latent activity be observable in the predicted firing rates. At latent dimensionalities (∼256D) necessary for comparable reconstruction performance to DynISys, approximately 30% of the total latent variance of the MR-LFADS model lay in the readout null space (Extended Data Fig. 5). This activity does not correspond to any feature visible in the instantaneous predicted firing rates, and therefore reflects a model hallucination.

To examine how target information was represented over time, we visualized single-trial latent activity together with time-specific target-decoding planes and projections onto those planes (Fig. 3B). Notably, we found that the optimal decoding plane in PMd rotated over the course of the trial, possibly reflecting the storage of information in an M1 output-null subspace prior to transmission to M1 later in the trial (Kaufman et al. 2014).

Information about the upcoming target can be observed in PMd prior to M1, raising an obvious question: is PMd the *source* of this information in M1? We tested this using the same decoding procedure as in Fig. 3A, and found that ĉ_*PMd*−>*M*1_ became predictive of the upcoming target around the time of target entry, after PMd’s own latents 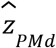 and before a corresponding rise in 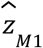 or ĉ _*M*1−>*PMd*_ (Fig. 3C). To highlight these relative timings, we visualized the target encoding in ĉ _*PMd*−>*M*1_ (Fig. 3D). Comparing Fig. 3 panels B and D illustrates a flow of target information through the model compartments around the time of current target entry. These results reveal that despite being fully unsupervised with respect to the behavioral task, DynISys infers inter-area communication signals consistent with the hypothesis that PMd transmits target information to M1 during motor preparation.

### DynISys infers input-dependent fixed points in M1 that encode upcoming reach

A primary hope for multi-compartment dynamics models is that they can be interrogated to understand how computations arise from the interactions between intrinsic dynamics and inputs – i.e., how the dynamics 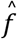 are functionally modulated by inputs ĉ and û (Fig. 4A). Here we tested whether DynISys could help reveal how M1’s intrinsic dynamics are modulated by inferred inputs during motor preparation. Prior work suggests that M1 contains a family of stable, attractive fixed points that encode the upcoming motor plan (Inagaki et al. 2019). We hypothesized a mechanism for motor preparation in which the fixed point expressed on any particular trial (Fig. 4B) is determined by inputs from PMd. This arrangement would imply a continuous manifold of potential fixed points that facilitates flexible selection of desired motor plans from a diverse repertoire (Fig. 4B, right). Such input-dependent fixed points could organize preparatory states that serve as initial conditions for subsequent movement-generating dynamics (Churchland et al. 2010, Shah et al. 2025).

**Figure 4.**
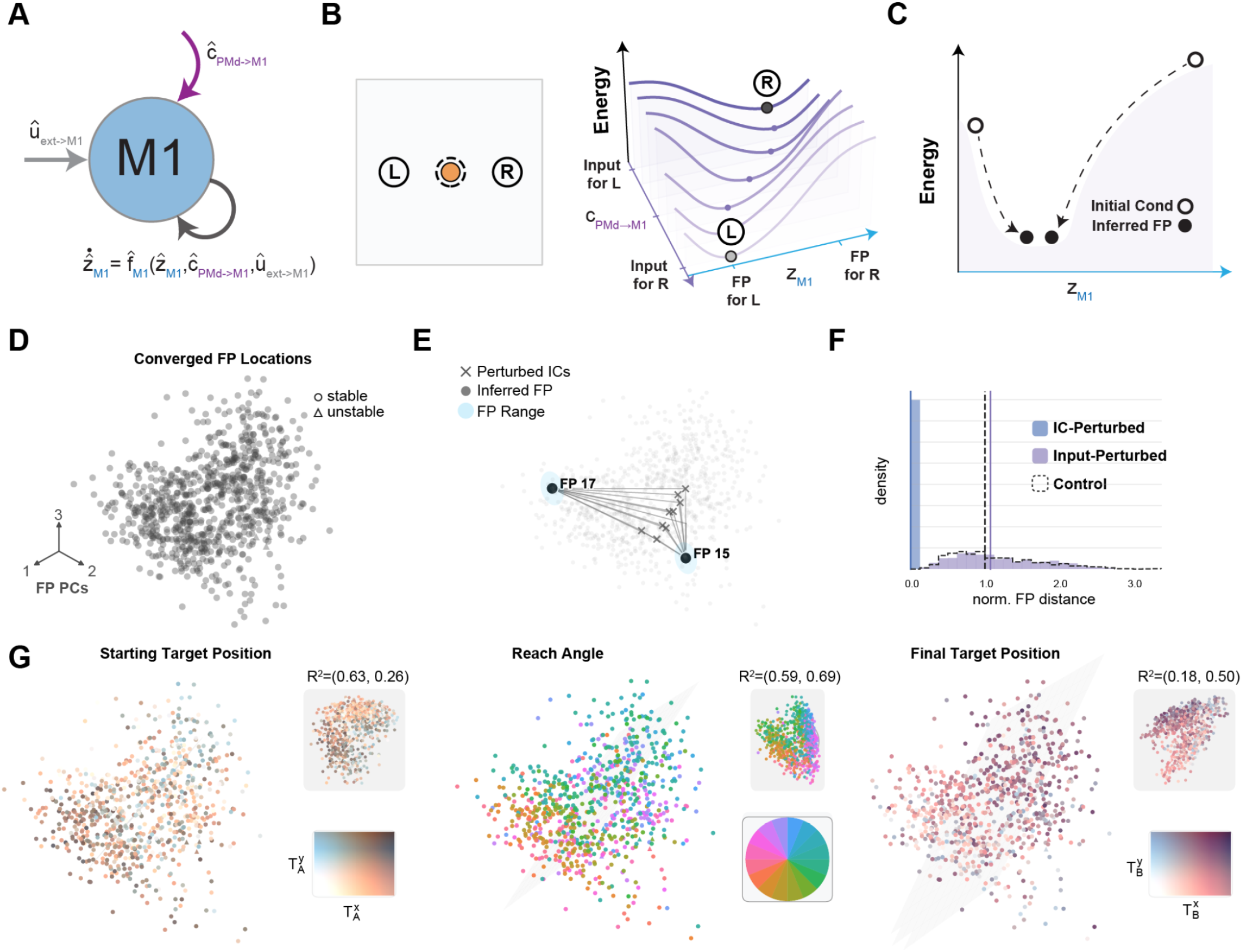
M1 dynamics exhibit input-modulated fixed points that encode features of the upcoming reach. **(A)** Schematic of inferred M1 dynamical compartment. M1 dynamics are a function of inferred external inputs and communication from PMd. **(B)** Schematic of input-modulated energetic landscape in M1 for two example reaches (L/R, orange circle indicates starting cursor location). Right: Input from PMd (purple) modulates the schematic energy landscape (vertical axis, black) along target-encoding preparatory dimensions in M1 (blue). When PMd input signals indicate reach L, the resulting M1 dynamics express a stable, attractive fixed point (FP) that encodes the motor plan for reach L. Intrinsic M1 dynamics drive M1 neural activity to that fixed point. **(C)** Example fixed point finding procedure. Random initial conditions were chosen in M1, and optimization seeks energetic local minima of dynamical landscape. **(D)** Top 3 PCs of inferred M1 fixed points from optimization with each trial’s own inputs and initial condition. 797/821 (>97%) of fixed point optimizations converged to locations with neural speed lower than 1e-6. Of these inferred fixed points, all were found to be stable (797/797, circles). **(E)** Example fixed point descents, computed using two trials’ inferred inputs (trials 15 & 17) and 10 randomly selected initial conditions. Converged fixed point locations are highlighted with cyan circles. Example fixed point optimizations tend to converge to locations organized by the external input, rather than initial conditions. **(F)** Histogram of latent distances from same-trial fixed point locations (panel D) to fixed points computed with the same trial’s input (cyan) or initial condition (purple). Shuffled control (distance from original fixed point to fixed point inferred by random trial’s IC and input) shown as dashed line. Static input fixed points tend to be much closer to the same-trial fixed point than static IC fixed points. **(G)** Inferred fixed point locations encode features of upcoming reach. Each column shows same-trial fixed point locations color coded by features of upcoming reach. Colormap of feature and decoder output / performance shown as insets in each column (bottom, top insets, respectively). Left: Starting target position. Middle: Reach angle. Right: Final target position.

We first searched for M1 fixed points under each trial’s original inferred inputs. Each search began at the latent state *z*_0_ inferred 200 ms after entry into the preceding target and minimized latent-state speed while holding the corresponding inputs {*c, u*} constant (Golub and Sussillo 2018; Driscoll et al. 2024; Fig. 4C; Methods). We call the resulting baseline solution the trial’s *unperturbed M1 fixed point*, against which we compare subsequent searches with altered starting states or inputs. Searches converged to slow states (median q = 8.4 × 10^−12^, 95% CI: [7.7, 9.1] × 10^−12^). Of 821 solutions, 797 met the fixed-point criterion (q < 10^−6^); all 797 were locally stable, with strictly negative Jacobian eigenvalue real parts (Fig. 4D).

We next asked if the inferred fixed points were purely a property of the inputs that modulate M1’s dynamics, or if they were also determined by initial states. We re-ran the fixed point finding procedure many times for each trial, replacing either 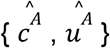 or 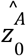 for a given trial A with the corresponding value from a different randomly chosen trial α. *IC-perturbed* searches used the trial’s own inferred inputs 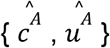 but another trial’s initial condition 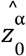, and *input-perturbed* searches used the trial’s own initial condition 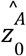 another trial’s inferred inputs 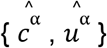. We computed the latent distance from each trial’s unperturbed fixed point to these perturbed fixed points to quantify the relative importance of starting state and input to the recovered location of the inferred fixed point.

We visualized IC-perturbed fixed point finding trajectories for two example trials (Fig. 4E) and found that fixed point descents all converged to a point very close to that trial’s unperturbed fixed point location. This trend was seen across trials and repeats: IC-perturbed fixed points remained close to their unperturbed counterparts, whereas input-perturbed fixed points fell far away, approaching the distance expected when both ingredients are drawn from a random trial (Fig. 4F). These results show that inputs, not initial conditions, determine the inferred M1 fixed point, suggesting that M1 dynamics have strong input-dependent modulation.

Finally, we asked whether fixed point location carries information about the upcoming movement. We found that unperturbed fixed points varied systematically with the location of the starting target and with the location of the upcoming target. We fit linear decoders from fixed point location to each of these features and found that they provided substantial target information (Fig. 4G, mean R^2^ values of 0.45, 0.64, 0.34 for starting position, reach angle, final target position, respectively), consistent with M1 dynamics using fixed points to store features of an upcoming reach. We found similar results in Monkey M (Extended Data Fig. 6).

### In-silico perturbations predict input-driven dynamical mechanisms of movement preparation and initiation

A major appeal of input-driven nonlinear dynamics models such as DynISys is that they enable simulated experiments that highlight causal relationships inferred by the model. In this section, we use targeted input perturbations to help interpret how M1’s inputs and intrinsic dynamics interact during movement planning and initiation.

First, we sought to understand *how* inputs from PMd modulate M1 preparatory dynamics to create target-encoding fixed points. Using the previously described linear decoders from ĉ_*PMd*−>*M*1_ to upcoming target location, we identified the target-encoding directions in the PMd→M1 input space (Fig. 5A, purple vectors). We injected perturbations along chosen directions in that target-encoding plane by adding them to each trial’s inferred PMd→M1 input ĉ_*PMd*−>*M*1_. We then recomputed M1’s fixed points under the perturbed input (see Methods) and compared the location of the perturbed fixed point to the original unperturbed fixed point.

**Figure 5.**
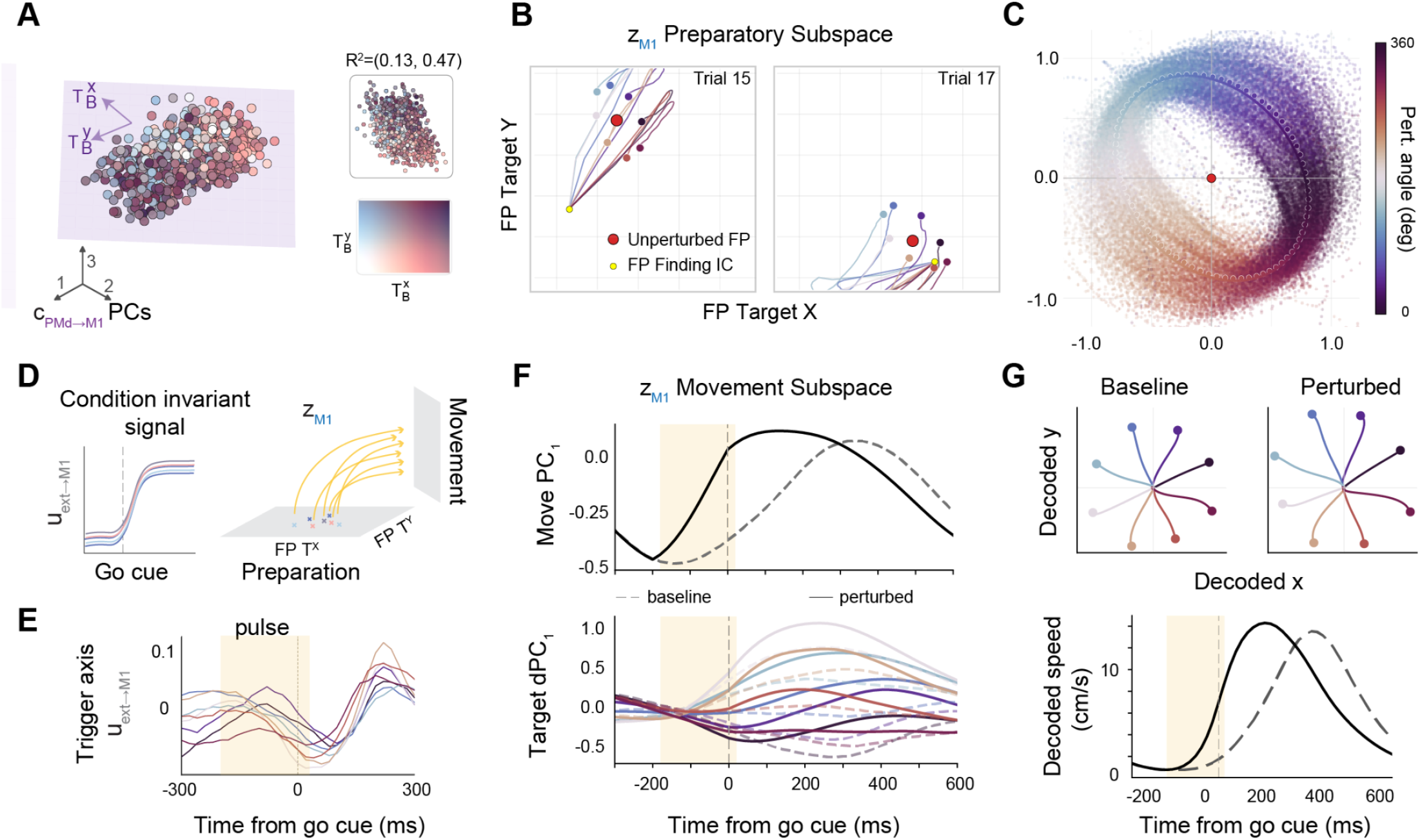
Simulated perturbations differentiate functional roles of inferred PMd and external inputs to M1. **(A)** PMd→M1 communication channel activity ĉ_*PMd*−>*M*1_ projected onto its top 3 PCs, colored by upcoming reach target (X/Y, see inset). A linear decoder (purple plane) maps this channel onto the next target; inset shows decoded vs. true target performance R^2^. **(B)** For two example trials, converged M1 fixed points obtained by descending the flow field under systematically perturbed PMd→M1 inputs (colored by perturbation direction), decoded into target space. Red marker = unperturbed fixed point; yellow marker = FP finding initial condition. **(C)** Across all trials, fixed point displacement (perturbed − unperturbed, decoded target space) as a function of perturbation direction (color, 0-360°), with per-direction means and best-fit response ellipse. M1 fixed points shift in the direction predicted by the communication channel’s decoded target. **(D)** Schematic of the causal hypothesis tested in E–G. A condition-invariant trigger in the ext→M1 channel after the go cue propagates to the downstream M1 circuit, initiating movement away from the preparatory state. **(E)** Condition-averaged ext→M1 activity colored by relative reach direction and projected onto the identified trigger axis (shaded window, pulse epoch). **(F)** The pulse perturbation shifts M1 dynamics earlier in time within the movement subspace, visualized by trial-averaged activity along the leading movement PC (top) and direction-conditioned activity along the leading target-dependent dPCA component (bottom) (dashed = baseline, solid = perturbed; black = grand average across directions). **(G)** Top: decoded relative hand trajectories (x, y) from M1 latents for baseline (left) and perturbed (right) trials for eight example trials; × marks the go cue. Perturbed trajectories move toward the planned target. Bottom: average decoded speed across trials aligned to go cue (dashed = baseline, solid = perturbed); the pulse shifts the speed profile earlier in time, indicating that the perturbation triggers premature movement.

We found that a perturbation in a given direction in the communication space tended to shift the inferred fixed point in the same direction in the fixed point target-decoding space (Fig. 5B). This effect was systematic across trials and monkeys. We applied a dense set of perturbation directions to each trial and recentered the resulting fixed-point locations on the unperturbed location. The shifts traced a ring whose orientation followed the perturbation’s target direction (Fig. 5C; Extended Data Fig. 7). This perturbation analysis elevates the correlative analysis of Figure 4 to a model-derived mechanistic hypothesis that can be tested experimentally: specific, target-encoding dimensions of the PMd→M1 inputs set the location of M1’s fixed points to encode features of the upcoming reach.

DynISys infers communication from recorded areas and external inputs from unobserved sources separately, allowing us to distinguish their relative contributions to modeled motor cortical dynamics. Previous work identified a condition-independent signal in PMd and M1 around movement onset (Kaufman et al., 2016), potentially driven by a midbrain–thalamocortical pathway that triggers movement initiation (Inagaki et al., 2022). We next asked whether DynISys identified such a “trigger” signal in the inferred external inputs that controls when M1 leaves the preparatory state and enters a movement-generating dynamical regime. We predicted that artificially advancing this signal would prematurely initiate the prepared movement, supporting a mechanistic model in which PMd→M1 communication specifies the movement to be performed while condition-invariant external inputs control when that movement is initiated (Fig. 5D).

To identify a candidate trigger axis, we used dPCA to find a condition-invariant component in the inferred ext→M1 channel û_*ext*−>*M*1_ that exhibited an abrupt change following the go cue (Fig. 5E, see Methods, ‘Movement initiation analyses’). We then performed an *in-silico* experiment to assess how a 200 ms square pulse injected along the trigger axis, starting 180 ms before the go cue, affected M1 latent activity over a window spanning −300 to 600 ms relative to the go cue. Perturbing the ext→M1 candidate trigger axis caused M1 population activity to exhibit movement-related activity substantially earlier than in baseline trials, as shown by projecting M1 latent activity onto its movement subspace (Fig. 5F). To quantify this shift, we estimated movement onset time using linear discriminant analysis and found that perturbation advanced onset by 182.2 ± 2.3 ms (mean ± SEM) relative to baseline (see *Methods*). This shift scaled with perturbation amplitude and timing (Extended Data Fig. 8) and was specific to the ext→M1 channel, as perturbing PMd→M1 input did not produce a comparable shift (Extended Data Fig. 9, blue trace).

We next asked whether this earlier transition corresponded to execution of a particular prepared movement rather than a nonspecific shift in M1 activity. We trained a recurrent neural network decoder to predict hand velocity from baseline M1 latent states over the interval from 400 ms before to 600 ms after the go cue. Applying this decoder to latent activity generated under the perturbed input produced movements that preserved the original movement directions (examples in Fig. 5G, top), but with speed profiles that were shifted earlier relative to the unperturbed trials (Fig. 5G, bottom). Thus, within the fitted model, stimulation along the putative ext→M1 trigger axis was sufficient to advance the transition from preparation to movement while preserving the direction of the prepared movement. Perturbations along the highest variance direction orthogonal to the trigger axis had mixed effects, with positive pulses producing a small onset shift and negative pulses advancing onset while also altering decoded movement direction (Extended Data Fig. 9).

Together, these in-silico perturbations suggest distinct functional roles for inputs to M1. During planning, inferred PMd→M1 communication determines the movement-specific preparatory state by controlling the location of M1’s input-dependent fixed point, while during initiation a condition-invariant component of the inferred external input controls when M1 leaves that state and enters movement-generating dynamics. This division provides a concrete, experimentally testable hypothesis for how inter-area and external inputs jointly organize movement preparation and initiation. More broadly, the analysis illustrates how multi-area dynamical models can serve as testbeds for formulating mechanistic hypotheses in the language of dynamics.

### DynISys aligns PMd and M1 latent dynamics across recording sessions into a common, behaviorally relevant latent space

Our ability to characterize the features reflected in neural population activity generally improves with the number of neurons observed (Kohn et al. 2016). However, the number of neurons that can be recorded simultaneously is limited by experimental constraints, so that any single session may only capture a partial view of the shared dynamics. Previous models have sought to stitch multiple datasets together to recover a single dynamical system that explains activity from many days of recordings, under the assumption that the same task engages the same dynamics across sessions (Pandarinath et al. 2018). However, for DynISys the problem of multi-session stitching is non-trivial, as the injective readout could restrict the model’s ability to learn a single dynamical rule that accounts for populations with widely variable firing statistics.

To overcome this issue, we developed a stitching strategy that enables DynISys to learn a single dynamical model from data pooled across sessions (Fig. 6A). Per-session, per-area session-alignment read-in matrices first project the variable-sized populations into a common input space; a single, shared DynISys core then infers the joint latent dynamics from these aligned inputs; finally, the latent is mapped back to neural activity through a shared injective readout together with per-session, per-area session-alignment read-out matrices that account for the varying number of units recorded in each session. The session-alignment read-in and read-out matrices are precomputed by a separate supervised alignment procedure and kept frozen during training of the DynISys core (see Methods).

**Figure 6.**
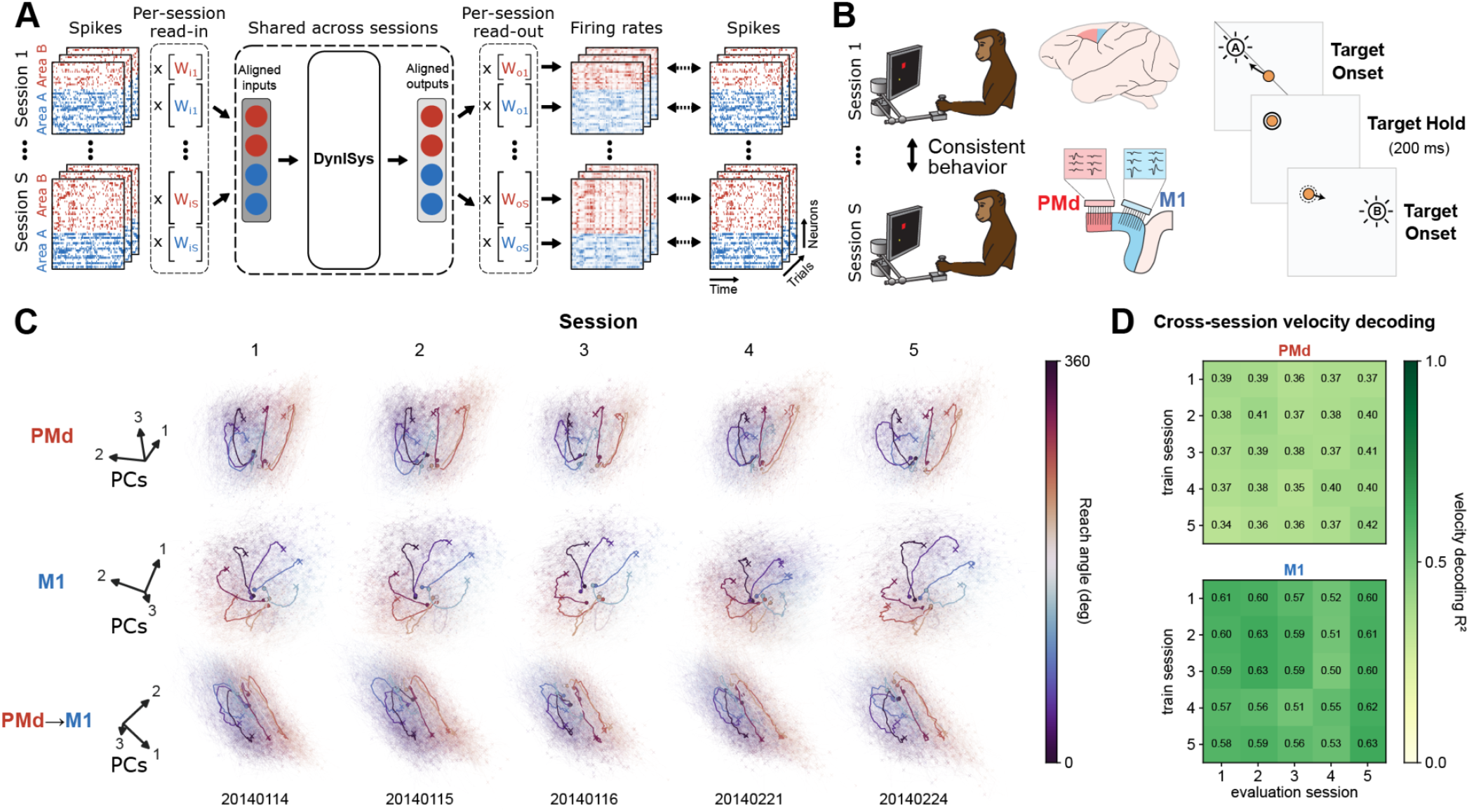
DynISys learns latent dynamics that are shared across multiple recording sessions. **(A)** Multi-session stitching strategy. For each session, spikes from areas A/B are projected by per-session, per-area session-alignment read-in matrices into a common input space of fixed dimensionality. A single DynISys core, shared across all sessions, infers the joint latent dynamics from these aligned inputs. The latent is then mapped back to each session’s neurons through an injective read-out followed by per-session, per-area session-alignment read-out matrices, yielding firing rate predictions that reconstruct the recorded spikes under the Poisson spiking model. Session-alignment read-in and read-out matrices were precomputed from individual areas and sessions (see Methods). **(B)** Behavioral task and recording setup. A monkey performed a random-target reaching task with consistent behavior across sessions. It made reaches to a series of individually presented targets, briefly holding on each target for ∼200 ms before the subsequent target was displayed. Spiking activity from PMd (red) and M1 (blue) were recorded simultaneously with two Utah arrays over five sessions spanning approximately six weeks (Glaser et al. 2018, Perich et al. 2025). **(C)** Single-trial latent trajectories (top-3 PCs) for PMd, M1, and PMd→M1 communication compartments (rows), shown separately for each of the five sessions (columns) and colored by reach angle (colorbar). We grouped reaches into eight octants based on reach directions and showed their averaged latent trajectories as thick lines; ∘ and × denote the starting and ending states, respectively. Although latent trajectories are inferred from and mapped back to a distinct set of neurons for each session, the trajectories share a common, direction-organized geometry across sessions. **(D)** Cross-session decoding of hand velocity. For each area, a linear decoder was trained on one session’s latents and evaluated on the latents of every session (within-session performance computed out-of-fold), yielding a 5 × 5 matrix of decoding accuracy (in R^2^ - rows represent training session and columns represent evaluation session). Cross-session (off-diagonal) accuracy is comparable to within-session (diagonal) accuracy, indicating that the inferred latents preserve a consistent mapping to behavior across sessions.

We applied this approach to a separate publicly available dataset in which a monkey performed a random-target reaching task over five sessions spanning approximately six weeks (Glaser et al. 2018). The dataset contains simultaneous spiking activity from PMd and M1 areas recorded using multielectrode arrays while the monkey was reaching to sequentially presented targets, briefly holding on each target (∼200 ms) before the next target appeared (Fig. 6B). We found that single-trial latent trajectories in PMd, M1, and PMd→M1 communication subspaces exhibited a consistent, directional structure that was preserved across all five sessions despite being inferred from distinct populations of neurons (Fig. 6C). We quantified this finding by training a linear decoder to predict a session’s hand velocity from the same session’s latents and evaluating its performance on latents inferred from other sessions. We found that decoders generalized well across sessions: cross-session (off-diagonal) accuracy was comparable to within-session (diagonal) accuracy in both PMd and M1, indicating that the relationship between the latent activity and behavior was preserved in the DynISys model (Fig. 6D). As expected, decoder performance was higher from M1 (R^2^ 0.5-0.63) than from PMd (R^2^ 0.34-0.42), in line with the more direct role of M1 in movement execution. The near-uniformity of each matrix confirmed that DynISys embedded all five sessions within a single, session-invariant latent space, the alignment that our stitching strategy was designed to achieve.

## Discussion

DynISys enables interpretation of functional computation from multi-area neurophysiological recordings by linking neural population models to analytical tools from nonlinear dynamical systems. This analytical toolset is mature: fixed-point finding and flow field characterization have been used for over a decade to explain how recurrent networks solve tasks, converting trained networks from opaque function approximators into systems whose computation can be described in the language of dynamics (Sussillo and Barak 2013, Golub et al. 2018, Maheswaranathan et al. 2019, Driscoll et al. 2024, Marraffa et al. 2026). However, the analytical toolset has been developed and validated on synthetic systems where ground truth is available by construction. Applying this toolset to models fit to neural recordings has been limited by the trustworthiness of the models themselves, as flexible models with unconstrained inputs can produce a variety of dynamical solutions with different computational properties (Kao 2019, Sedler et al. 2023, Versteeg et al. 2025). DynISys addresses this gap through three architectural features. Neural-ODE generators provide low-dimensional vector fields, injective readouts constrain latent states to remain recoverable from predicted firing rates, and low-dimensional communication channels constrain interactions between recorded areas. The result is a model that supports interrogation to test mechanistic hypotheses about the underlying circuit.

This work connects to a broader effort in NeuroAI to derive computational understanding from large-scale recordings. Much of that effort proceeds through representational description: encoding models and foundation models that predict neural responses to novel inputs, and probes, decoders, and alignment metrics that characterize what information a population carries and how it is geometrically organized (e.g., Wang et al. 2025). While impactful, these studies largely ask what variables are present in which neural populations, rather than specifying the mechanism by which those variables are used to generate useful output. We need trustworthy dynamical proxies of neural circuits that support simulated experiments, allowing researchers to perturb inferred states and inputs and formulate testable hypotheses about how their interactions generate behavior.

DynISys extends earlier approaches to modeling interacting neural populations. The communication subspace framework (Kaufman et al. 2014, Semedo et al. 2019) established that inter-area signaling can be confined to a small number of dimensions, recovered by regression between simultaneously recorded populations. CURBD (Perich et al. 2026) fits a recurrent network at the dimensionality of the recorded neurons and decomposes its currents into within- and across-area contributions, but learns no low-dimensional latent dynamics and does not separately infer inputs from unobserved areas. The multi-population recurrent switching linear dynamical system of Glaser et al. (2020) introduces per-population low-dimensional states, but approximates nonlinear interactions with a finite set of switching linear regimes and likewise infers no continuous external inputs. MR-LFADS (Liu et al. 2025) is the conceptually closest analogue to DynISys since both models jointly infer area-specific nonlinear dynamics, inter-area communication, and unobserved external inputs on single trials. MR-LFADS implements local dynamics with GRUs and maps generator states to rates through linear factor readouts, whereas DynISys uses neural-ODE dynamics with a readout explicitly constrained to be injective. Injectivity makes the model’s latent state recoverable from predicted rates, and thereby limits the model’s capacity to hallucinate structure in latent dimensions without affecting reconstructed activity. DynISys is consequently designed not only to identify communication, but to yield states, flow fields, and input-dependent fixed points that can be interpreted and perturbed.

Applied to PMd and M1, DynISys bridges previously separate accounts of motor preparation. First, Kaufman et al. (2014) showed that preparatory activity in PMd occupies dimensions with little influence on M1. DynISys infers such a channel directly from the recordings and assigns it a computational role: target-specific PMd→M1 input reshapes M1’s dynamical landscape to select the preparatory state toward which activity converges. Second, in contrast to the discrete attractor dynamics reported by Inagaki et al. (2019) in mouse anterior lateral motor cortex, our model infers a single, input-dependent fixed point per trial, drawn from a continuous manifold. While discrete attractors are a natural solution for a binary decision, they cannot flexibly specify the graded, continuously varying reaches of this task. Third, we show that this fixed point location sets initial conditions of subsequent movement-generating dynamics that can be evoked by an external trigger, in line with previous work showing that thalamic input triggers planned movements in M1 (Inagaki et al. 2022). Together, these findings support the hypothesis that inter-area communication configures downstream dynamics that store and ultimately generate the selected movement.

Although we demonstrate DynISys on intracortical recordings of the motor system, the approach is not restricted to these particular systems, brain areas, or recording modalities. The model is trained without access to task variables or behavior, and its application does not require strong constraints on experimental design. This raises the possibility of dynamical modeling in naturalistic behavioral tasks without well-defined conditions, and in other cognitive domains where computation is performed in internal states and where the relevant measurable variables may not be known *a priori*. In addition, the DynISys observation model is modular, allowing the Poisson link to be replaced to accommodate other recording modalities, with prior work suggesting that dynamical inference can transfer to two-photon calcium imaging (Zhu et al. 2022). Simultaneous multi-area recordings (multiple Neuropixels probes, chronic multi-array implants in humans, and large-scale two-photon imaging) are now routine in many laboratories, suggesting that we may soon enter a period where the primary bottleneck is no longer adequate datasets, but the ability to interpret them.

Several limitations motivate future work. The causal relationships identified here hold within DynISys and should be interpreted as experimentally testable hypotheses rather than established properties of the biological circuit. Their generality must also be evaluated across animals, tasks, and brain areas. Future pathway-specific perturbations could directly test the model’s central predictions: manipulating target-related PMd→M1 activity should alter the preparatory state selected in M1, whereas manipulating condition-invariant inputs should alter movement timing while preserving movement identity. Such experiments would help determine which aspects of the inferred mechanism are implemented in the brain.

## Methods

### Behavioral task and neural recordings

#### Kinarm Datasets

We analyzed simultaneous recordings from dorsal premotor cortex (PMd) and primary motor cortex (M1) of two monkeys performing a sequential reaching task with a Kinarm manipulandum (Michaels et al. 2025). On each trial the animal acquired a series of visual targets in sequence. Trials were of two types: on *Horizon-1* trials only the target currently being acquired was visible, whereas on *Horizon-2* trials the next, upcoming target was displayed at the same time, allowing the animal to plan the upcoming reach while completing the current one. After preprocessing, the analyzed dataset comprised the PMd and M1 single units listed in Table 1. Unless otherwise noted, analyses were restricted to Horizon-2 trials and aligned to entry of the current target; the times of target changes, and the identities of the current and upcoming targets at each moment, were computed from the recorded target positions (data/Kinarm/target_epoch_utils.py).

#### Mihili Datasets

For the multi-session analyses we used publicly available recordings from monkey Mihili performing a random-target reaching task (Glaser et al., 2018, Perich et al. 2025). In this task, the monkey moved a cursor with a planar manipulandum to acquire targets. Each trial consisted of four sequentially presented targets that appeared at random locations 5-15 cm from the previous target. After the cursor entered the target and held for approximately 200 ms, the next target was presented and the monkey was free to begin the subsequent reach without any imposed delay period. Spiking activity was recorded simultaneously from PMd and M1 with two 96-channel Utah arrays and sorted offline (Glaser et al., 2018). We analyzed five sessions spanning approximately six weeks (Table 2).

**Table 2:** Mihili dataset and preprocessing parameters. Per-session values are listed in chronological order (20140114, 20140115, 20140116, 20140221, and 20140224).

| **Parameter** | **Value** |
|---|---|
| Spike bin width | 20 ms |
| Window (initial-condition + modeled segment) | 75 + 75 bins (3,000 ms); stride 25 bins |
| Cross-talk criterion | \|*r*\| > 0.025 between 2-ms spike counts |
| Task disengagement | gap > 2 s between consecutive task events |
| Reach inclusion | rewarded trials; displacement ≥ 2 cm; duration ≤ 1.4 s |
| Minimum mean firing rate | 1 Hz (−100 to +500 ms around target onset) |
| Held-in / held-out split | 80% / 20% |
| M1 units (total / held-in) | 47/38, 55/44, 58/46, 29/23, 31/25 |
| PMd units (total / held-in) | 73/58, 64/51, 73/58, 70/56, 69/55 |

### Preprocessing

#### Kinarm Datasets

Spike trains were binned and modeled over a fixed trial window. Single units were retained according to a recording-stability criterion and a minimum mean firing-rate threshold; we refer to the resulting inclusion set as the *Stable* dataset. For modeling, the neurons in each area were partitioned into held-in and held-out sets: only the held-in neurons were provided to the model’s encoder, while all neurons were reconstructed, so that held-out neurons measured the model’s ability to generalize to neurons that were not visible to the encoders. We binned the neural activity at 20 ms and sliced the data into overlapping 1600 ms slices (80 bins) with a stride of 400 ms (20 bins, 60-bin overlap between slices). The binned spikes, the held-in/held-out partition, and the aligned behavioral variables were written to HDF5 datasets consumed by the modeling pipeline. Bin width, trial window, inclusion threshold, split, and unit counts are given in Table 1; preprocessing code is under data/Kinarm/preprocessing/.

#### Mihili Datasets

Spike trains were binned and segmented into overlapping windows, each comprising an initial-condition segment, from which the encoder inferred the initial state, followed by the modeled segment. Units were retained according to a cross-talk criterion, which removed units with near-synchronous spiking on another channel, and a minimum mean firing-rate threshold. Of the six available sessions, one session (20140214) was excluded because it yielded only 22 sorted M1 units (32–70 in the other sessions), too few to estimate its M1 read-in and read-out reliably. As for the Kinarm dataset, units in each area were partitioned into held-in and held-out sets. Windows spanning periods of task disengagement were discarded, and contiguous blocks of each recording were reserved for validation. Reaches were defined by target onsets in rewarded trials, and their directions, given by the net cursor displacement, were binned into eight 45° sectors.

Preprocessing parameters and unit counts are given in Table 2.

### The DynISys model

DynISys models the recorded areas as a set of coupled latent dynamical systems, one compartment per area, together with inferred inputs that account for drive originating outside the recorded brain areas (implementation in ctd/data_modeling/models/LFADS/dynisys.py). Each compartment *a* carries a latent state 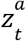; the architecture hyperparameters are listed in Table 3.

**Table 3:** DynISys architecture hyperparameters.

| **Parameter/Monkey** | **Monkey P** | **Monkey M** |
|---|---|---|
| Areas modeled | PMd, M1 | PMd, M1 |
| Per-area latent dimension *d* | 12 | 8 |
| Controller compartment dimension *d*<sub>*c*</sub> | 128 | 128 |
| Inter-area communication dimension | 3 (each direction) | 3 (each direction) |
| Controller→compartment input dimension | 3 | 3 |
| Generator MLP *f*<sup>*a*</sup> | 3 hidden layers × 128, ReLU | 3 hidden layers × 128, ReLU |
| Euler integration step ß | 0.1 | 0.1 |
| Encoder hidden size | 256 | 256 |
| Readout *g*<sup>*a*</sup> | injective flow (Stiefel + volume-preserving initialization) | injective flow (Stiefel + volume-preserving initialization) |
| Observation model | Poisson (smooth-ReLU link) | Poisson (smooth-ReLU link) |
| Fisher penalty scale (minSV_scale) | 1e-3 | 1e-4 |
| Fisher SNR target *s*<sup>⋆</sup> | 2 | 2 |

#### Generative model

Let 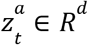 denote the latent state of compartment *a* at time *t* and 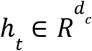 for the state of a shared controller compartment (dimensions in Table 3). Compartments interact only through explicit, low-dimensional stochastic channels: each ordered pair of compartments carries a communication signal, and the controller supplies each compartment with an inferred input,

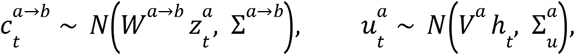

so that all inter-area influence is confined to a few interpretable dimensions. Collecting the signals that reach compartment *a* into 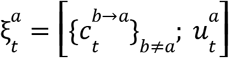, each compartment’s state evolves under its own generator *f*^*a*^—a continuous-time vector field parameterized by a multilayer perceptron (Table 3)—integrated with a fixed-step (Euler) update of size ß at the bin resolution,

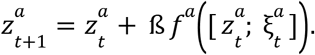

This neural-ODE parameterization decouples the expressivity of each compartment’s dynamics from the dimensionality of its latent state, and confining all inter-area influence to the low-dimensional channels 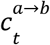 keeps the communication between areas easy to interpret.

#### Readout

Each compartment’s latent state is mapped to firing rates by an injective normalizing-flow readout (InjectiveFlowReadout) followed by a positive rectifying activation. Specifically, the readout is composed of a column-orthogonal (Stiefel) embedding from the latent dimension into the neural dimension followed by a sequence of flow steps initialized to be volume-preserving (orthogonal permutations interleaved with additive coupling layers), 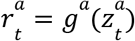 . A smooth rectifying link ρ (Smooth-ReLU) then produces firing rates, assuming Poisson spike counts,

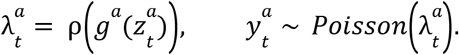

Because *g*^*a*^ is injective, the latent state remains recoverable from the rates, which guards against the model placing structure in directions the rates cannot express.

#### Inference

Latent states, communication signals, and inputs were inferred by variational bidirectional recurrent encoders: one pathway produced each compartment’s initial-condition posterior (a diagonal Gaussian, 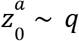 and another the time-varying input posterior, while the channel signals 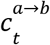 and 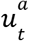 are sampled from the Gaussian posteriors in the rollout above. The canonical dataset used a causal encoding window for the initial condition, so that the inferred initial state cannot draw on future activity—in contrast to the non-causal IntLFADS baseline (Training).

#### Restitching

To recover continuous trajectories from windowed inference, we reconstructed each recording segment from the overlapping 1.6-s snippets (80 bins at 20 ms resolution), offset by 400 ms (20 bins). Each snippet was inferred independently and placed at its original temporal position. At each time bin, we averaged the available estimates of compartment states, communication signals, inferred inputs, and firing rates across overlapping snippets, giving each contributing estimate equal weight. To reduce initialization transients, we excluded the first five bins (100 ms) of each snippet except the first snippet in a segment. Behavioral variables and task events were aligned to the same reconstructed time base for subsequent analyses. Restitching was performed separately within each contiguous segment of the recording.

#### Objective

The model was trained by minimizing the negative evidence lower bound together with two regularizers,

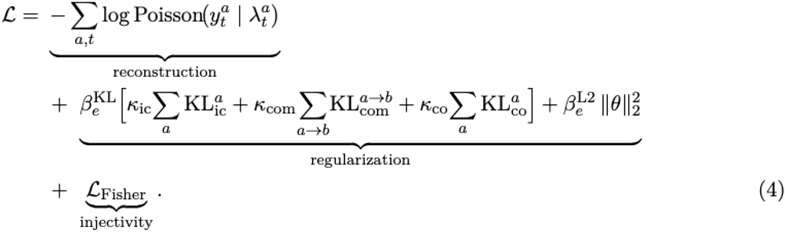

The three Kullback–Leibler terms penalize, respectively, each area’s initial-condition posterior against an *N*(0, *I*) prior, each inter-area communication signal 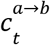 against its prior, and each controller→area input 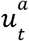 against its prior (all priors static-mean Gaussians); 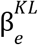 and 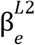 ramp the penalties linearly from zero over a fixed number of warmup epochs. Each of these terms can be scaled independently by κ_*ic*_, κ_*com*_, and κ_*co*_ (kl_ic_scale, kl_com_scale, kl_co_scale) to regularize the initial conditions, communication signals, and controller inputs toward their priors. In the canonical model reported here only the controller-input penalty is active (κ_*co*_ > 0); the initial-condition and communication penalties are disabled (κ_*ic*_ = κ_*com*_ = 0). These values, and the warmup schedule, are given in Table 4.

**Table 4:** Training hyperparameters for the canonical model.

| **Parameter** | **Value** |
|---|---|
| Optimizer | AdamW (β<sub>1</sub> = 0.9, β<sub>2</sub> = 0.999, ϵ = 10<sup>−8</sup>) |
| Weight decay | 0 |
| Initial / minimum learning rate | 1e-3, 1e-5 |
| LR scheduler | ReduceLROnPlateau (factor 0.95, patience 6) |
| Batch size | 256 |
| Maximum epochs | 500 |
| KL / L2 warmup | 0–40 epochs (linear) |
| Input prior (co_prior_type) | MultivariateNormalStaticMean |
| KL scale, controller inputs κ<sub>*co*</sub> | 10<sup>−3</sup> |
| KL scale, communication κ<sub>*com*</sub> | 0 |
| KL scale, initial conditions κ<sub>*ic*</sub> | 0 |

#### Readout-identifiability penalty

The injective flow *g*^*a*^ makes the latent→rate *embedding* one-to-one, but the readout must also pass through a rectifying link ρ to produce non-negative firing rates, and that link can reintroduce blind spots. Where it sits in its flat regime—neurons at or near zero rate, where ρ ^′^ ≈ 0—the composed map ρ ∘ *g*^*a*^ can become poorly conditioned, as there can be latent directions that barely move the firing rates even though the flow itself is invertible. Injectivity of the readout therefore does not on its own guarantee that every latent direction produces changes in firing rates that are visible above the noise level of the Poisson neural activity. To close this gap, a final term *L*_*Fisher*_ pressures the latent state to be *locally* recoverable from firing rates (FisherScaledJacobianMinSingularLossGram; scale in Table 3). At sampled latent states we form the Jacobian of the readout composed with the link function, *J*^*a*^ = ∂λ^*a*^ /∂*z*^*a*^, and scale it by the Poisson Fisher metric, 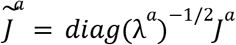, so that its singular values measure how distinguishable a latent perturbation is relative to spiking (Poisson) noise. A soft hinge on the smallest singular value,

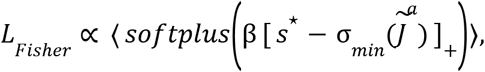

with ⟨ · ⟩ averaged over sampled states and compartments, pushes the smallest Fisher-scaled singular value above the SNR target *s*^⋆^ (Table 3).

Intuitively, this is a signal-to-noise floor on the readout. Under the Poisson model a neuron firing at rate λ has spike-count fluctuations of standard deviation 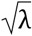, so the Fisher scaling *diag*(λ^*a*^) ^−1/2^ re-expresses how much a latent perturbation moves each firing rate in units of that neuron’s own noise; the singular values of 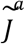 are then the population signal-to-noise ratio per unit of latent displacement, and σ_*min*_ is the value for the *least*-observable direction. Setting *s*^⋆^ = 1 asks that even this worst direction move the population rates by at least as much as the spiking noise: a direction with σ_*min*_ < 1 would change the rates by less than the trial-to-trial Poisson fluctuations, and so be effectively buried in the noise, present in the latents but unrecoverable from a single trial’s spikes. The penalty thus complements the *global* injectivity of the flow readout, which keeps the latent state formally invertible from noise-free rates, with a *local* pressure that no latent direction is flattened by the rectifying link or washed out by spiking noise.

### DynISys training

DynISys models were trained with the AdamW optimizer under a ReduceLROnPlateau learning-rate schedule that monitors a smoothed validation reconstruction loss, with early stopping on the same metric and the KL and L2 penalties ramped linearly over an initial warmup. The optimizer, schedule, batch size, epoch budget, input prior, and KL scales are listed in Table 4. Hyperparameters were swept with Ray Tune under Hydra configuration management, and training runs were logged with Weights & Biases.

### Single-neuron PSTH analyses

Peri-stimulus time histograms (PSTHs) were computed separately for PMd and M1 using Horizon-2 events pooled across the training and validation data. Observed spike counts were converted to firing rates and smoothed along the reconstructed recording with a normalized, one-sided Gaussian kernel (σ = 20 ms, truncated at 3σ), using only the current and preceding bins. DynISys-inferred firing rates were analyzed without additional smoothing. Activity was extracted in 600-ms windows aligned to three task events: the appearance of upcoming target B (−200 to +400 ms), entry into current target A (−300 to +300 ms), and the appearance of target C, when B became the current target (−400 to +200 ms). Trials lacking complete data over the analysis window were excluded.

For each alignment, events were grouped according to the direction of the vector from the hand position at the alignment event to target B. Thus, the grouping referred to the upcoming target for the first two alignments and the current target for the third alignment. Directions were divided into 16 equally spaced angular bins (22.5° each). For each neuron, PSTHs were calculated by averaging activity across events within each direction bin; bins containing fewer than three events were omitted. The same alignment and grouping procedures were applied to observed and DynISys-inferred activity.

To reduce modulation associated with the position of target A, we additionally computed residualized PSTHs for the target-A-entry and target-C-onset alignments (Fig. 2E). For each alignment, neuron, and time bin, we used ordinary least squares to predict activity from target A’s absolute horizontal and vertical coordinates, including an intercept:

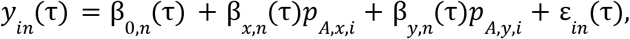

where *i* indexes events, *n* indexes neurons, and τ denotes time relative to alignment. Target A was the current target at its entry and the previous target at target-C onset. Regressions were fitted across events before direction grouping, separately for smoothed observed activity and DynISys-inferred rates. We subtracted the complete fitted prediction, including the intercept, from each event’s activity and recomputed direction-grouped means and standard errors from the residuals. This procedure removes the component linearly explained by target A’s position within the analyzed sample.

PSTH agreement was quantified by concatenating the observed and inferred condition means across time, retained direction bins, and included task phases. We calculated *R*^2^ separately for each neuron, using the smoothed observed PSTHs as the reference, and averaged these values across neurons within each area.

### Baseline Models

We summarize important differences between DynISys, IntLFADS, and MR-LFADS in Extended Data Table 1.

#### IntLFADS

As a baseline for Figure 3A, we trained IntLFADS, which shares DynISys’s multi-compartment structure but replaces the neural-ODE generators with standard recurrent (GRU) generators and the injective-flow readout with a linear readout, and, as in standard LFADS, infers both its initial conditions and its inputs from a non-causal (bidirectional) encoder. IntLFADS was trained on a dataset variant (NonCausalICs) in which the encoder window extends across the trial, so that its inferred initial condition can draw on future activity.

#### MR-LFADS

We trained MR-LFADS models using the authors’ implementation (Liu et al. 2025, https://github.com/golub-lab/MR-LFADS), with the same preprocessed recordings, held-in neurons, and training–validation partitions used for DynISys. Initial conditions were inferred from the preceding activity window. Communication was computed from reconstructed firing rates, capped at 20 expected spikes per bin for numerical stability. Separate predictors reconstructed held-out neurons from detached latent factors, preventing their prediction losses from updating the latent dynamics. Model and training hyperparameters are listed in Table 5. Comparisons used final training checkpoints and posterior-mean inference, with performance summarized as mean ± SEM across seeds.

**Table 5:** Hyperparameters for MR-LFADS models.

| **Parameter** | **Value** |
|---|---|
| Per-area latent/factor dimension *d* | [5, 8, 12, 16, 24, 32, 64, 128, 256] |
| Seeds per dimensionality | 5 |
| Inter-area communication dimension | 8 |
| External communication dimension | 8 |
| Per-area Controller compartment dimension *d*<sub>*c*</sub> | 128 |
| Input encoder delay | 1 bin |
| Priors on initial conditions, inferred inputs, and communication | *N*(0, 0. 1*I*) |
| KL scale, controller inputs κ<sub>*co*</sub> | 1e-1 |
| KL scale, communication κ<sub>*com*</sub> | 1e-1 |
| KL scale, initial conditions κ<sub>*ic*</sub> | 1.0 |
| IC and external inputs KL warmup | 50-250 epochs (linear) |
| Communication KL warmup | 150-250 epochs (linear) |
| Recurrent L2 weight | 1e4 (normalized by # parameters) |
| L2 warmup | 0-80 epochs (linear) |
| Initial / minimum learning rate | 4e-3, 1e-5 |
| LR scheduler | ReduceLROnPlateau (factor 0.95, patience 6) |
| Batch size | 128 |
| Gradient Norm clipping | 200 |
| Recurrent state clipping | 5 |
| Maximum epochs | 350 |
| Optimizer | AdamW (β<sub>1</sub> = 0. 9, β<sub>2</sub> = 0. 999, ϵ = 10<sup>−8</sup>) |
| Early stopping | Smoothed recon loss, patience 50 epochs (after warmups complete) |

#### CURBD

We fit current-based recurrent neural networks to the monkey P dataset using the official Rajan Lab CURBD implementation (Perich et al. 2026, https://github.com/rajanlab/CURBD). Networks contained one recurrent unit per recorded neuron, comprising 63 M1 and 64 PMd units. Spike counts were Gaussian-smoothed (σ = 40 ms, acausal) and converted to firing rates. Recurrent weights were trained using the implementation’s FORCE recursive least-squares procedure, with training and validation data separated by contiguous recording groups. We evaluated frozen-weight predictions against observed firing rates, testing both autonomous rollouts initialized once per group and reconstructions with neural-state resets at go cues. Across our models, we found that they quickly learned to estimate differences in mean firing rates across neurons but beyond that captured little of their temporal modulation.

To test whether the fitting procedure could reproduce neural activity in a more restricted setting, we trained the official implementation on a single eight-second segment, while also supplying resets at each go-cue in the short snippet. After 100 training passes, the model explained significant temporal variance in firing rates, measured relative to each neuron’s mean, when go-cue resets and the fixed training-noise realization were retained. These resets explicitly supervised the network by supplying observed neural population states throughout the trajectory; even with this additional supervision, the broader model explained very little temporal variation in firing-rates for held-out segments.

### Fixed point and communication analyses

#### Fixed points of the M1 generator

We located fixed points of M1’s learned vector field with a batched gradient-based search (damped Gauss-Newton). Starting from many random initial conditions and holding each trial’s inferred input fixed, we minimized the squared velocity ‖*f*^*M*1^(*z, c, u*)‖^2^ while holding the inferred communication and external inputs constant (optimizer and iteration settings in Table 6). At each recovered fixed point we evaluated the Jacobian of the generator and classified the point as stable if all of its eigenvalues had a negative real part.

**Table 6:**
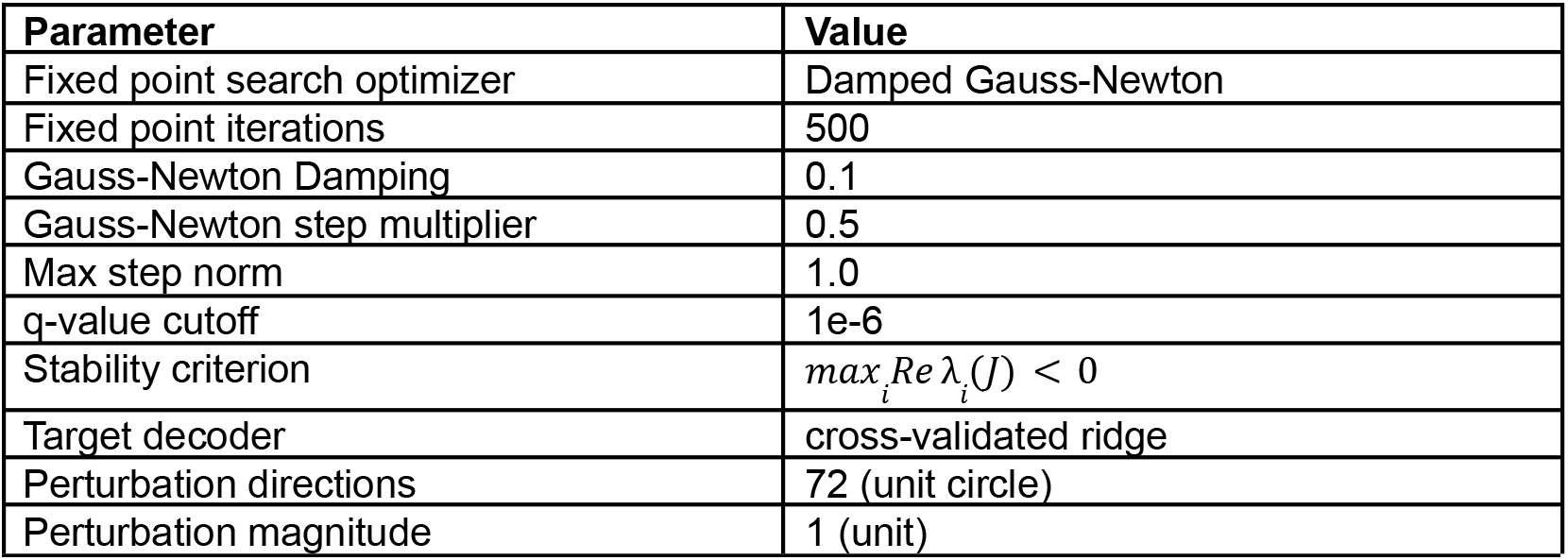
Fixed point and communication analysis parameters.

| Parameter | Value |
|---|---|
| Fixed point search optimizer | Damped Gauss-Newton |
| Fixed point iterations | 500 |
| Gauss-Newton Damping | 0.1 |
| Gauss-Newton step multiplier | 0.5 |
| Max step norm | 1.0 |
| q-value cutoff | 1e-6 |
| Stability criterion | max<sub>i</sub>Re λ<sub>i</sub>(J) < 0 |
| Target decoder | cross-validated ridge |
| Perturbation directions | 72 (unit circle) |
| Perturbation magnitude | 1 (unit) |

#### Input versus initial-condition controls

To test whether the inferred input or the initial condition determines the converged fixed point, we repeated the search while (i) holding the input fixed and drawing initial conditions from other trials, and (ii) holding the initial condition fixed and substituting other trials’ inputs. For each fixed point we decoded the upcoming-target position with a linear decoder fit to fixed point location and measured the error against the held target, comparing both conditions to a trial-shuffled chance distribution.

#### Communication decoding

The inferred PMd→M1 and M1→PMd channels were read directly from the fitted model. We decoded the upcoming target from each group of latents as a function of time using cross-validated linear (ridge) decoders aligned to entry of the current target, reporting held-out *R*^2^.

#### In-silico perturbation

To probe the input–landscape coupling causally within the model, we perturbed the inferred PMd→M1 communication along directions in its two-dimensional target-decoding plane (a dense set of unit-circle directions; Table 6), recomputed M1’s fixed points under the perturbed input, and measured the resulting shift of each fixed point relative to its unperturbed location.

### Movement initiation analyses

#### dPCA trigger-axis identification

We identified a condition-invariant trigger axis in the inferred external input to M1 (ext→M1) using demixed principal component analysis (dPCA). We aligned ext→M1 activity to the go cue, grouped all trials into eight upcoming reach direction conditions, and averaged activity within each condition over a 0–300 ms post-cue window. dPCA separated variance associated with upcoming direction, time, and their interaction; we defined the trigger axis as the leading decoder axis of the time marginalization and normalized it to unit length. Because the sign of a dPCA axis is arbitrary, we chose the orientation of the axis that made its condition-averaged projection more positive in the second half than in the first half of the 0–300 ms analysis window.

#### In-silico trigger perturbation

To test whether activity along this axis could initiate the prepared movement, we added a positive boxcar pulse to the ext→M1 input 180 ms before the go cue. The pulse lasted 200 ms and had an amplitude equal to 5 times the standard deviation of the trigger-axis projection, estimated from the first 200 ms after the go cue. We then reran the M1 generator while leaving the remaining inferred inputs and initial conditions unchanged.

#### Movement subspace identification

To visualize perturbation-evoked movement-related M1 dynamics separately from preparatory activity, we first estimated an M1 preparatory subspace from unperturbed H2 trials. Preparatory activity was sampled during a 200-ms window beginning 250 ms after the preceding target entry; trials were included only when the complete window ended before the upcoming go cue. PCA was applied to the pooled M1 latent activity from these windows, and the smallest set of components explaining at least 90% of the preparatory variance was removed. We then fit a second PCA to the residualized, movement-onset aligned M1 activity from −50 to 250 ms relative to movement onset to identify the movement related subspace. To visualize target-dependent dynamics within this movement subspace, we fit dPCA to unperturbed activity from 0 to 300 ms after the go cue, combining target and target-by-time marginalizations, and projected both unperturbed and perturbed activity onto the leading target-dependent component.

#### Movement onset detection

We identified a movement-related M1 dimension using linear discriminant analysis (LDA). The classifier was trained on movement-onset-aligned recurrent M1 latent activity to distinguish a quiescent epoch (−300 to −100 ms relative to behavioral movement onset) from a movement epoch (0 to 200 ms). The resulting affine decision score was used as a one-dimensional movement detector. For each trial, neural movement onset was defined as the first upward crossing of the LDA decision boundary that remained above threshold for at least three consecutive 20-ms bins (60 ms). Crossings were searched between −300 and 600 ms relative to the go cue.

#### Movement decoding

We trained a GRU decoder to predict two-dimensional hand velocity from the unperturbed go cue aligned M1 latent trajectories. The decoder was trained using all trials, with 15% held out for validation and early stopping. We applied the same decoder to the baseline and perturbed M1 trajectories, integrated the decoded velocity to obtain position, and compared the resulting movement direction and speed. Example trials were selected randomly with a fixed seed; population summaries included all trials. Figure-generation code for these analyses is in data/Kinarm/Figure_Generation/FigureExecution/.

### Multi-session DynISys

#### Stitching approach

To fit a single model to sessions with different numbers and identities of units, we adapted a previous approach for multi-session stitching (Pandarinath et al. 2018). Each session *s* and area *a* had its own session-alignment read-in and read-out matrices and bias vectors, with all other components (the encoders, generators, controller, communication channels and the flow readout *g*^*a*^) shared across sessions. The session-alignment read-in projected the held-in spike counts *y*^*a*^_*s,t*_ onto an input space of fixed dimensionality, and the session-alignment read-out mapped the flow output, also of fixed dimensionality, onto the firing rates of the session’s units,

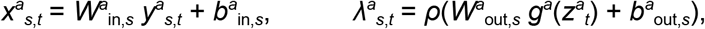

where *ρ* denotes the rectifying link (Smooth-ReLU). The session-alignment read-in and read-out matrices and biases were precomputed and frozen during training so that only the shared components were learned (see dimensions in Table 7).

**Table 7:**
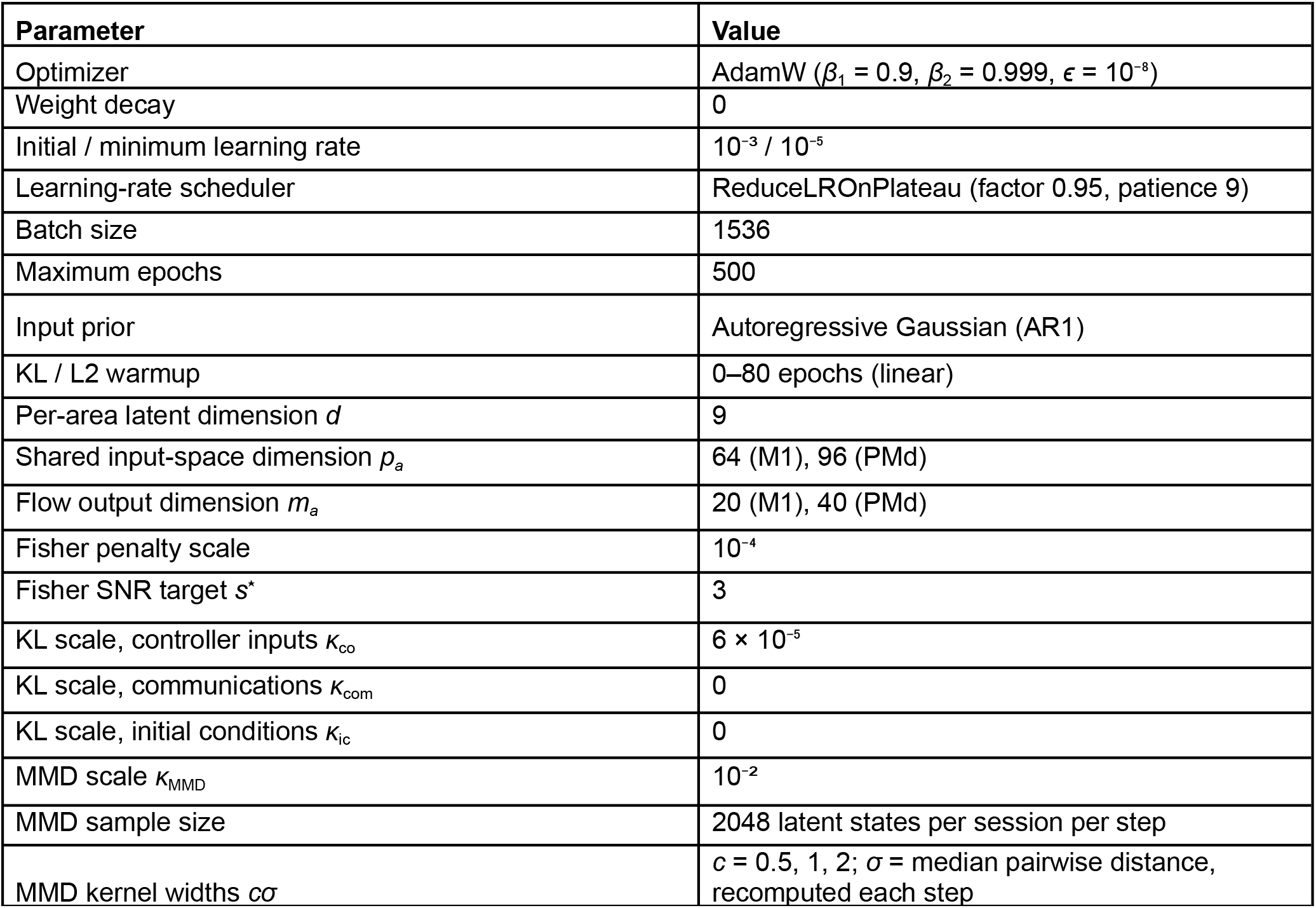
Multi-session model and training parameters.

| **Parameter** | **Value** |
|---|---|
| Optimizer | AdamW (*β*<sub>1</sub> = 0.9, *β*<sub>2</sub> = 0.999, *ϵ* = 10<sup>−8</sup>) |
| Weight decay | 0 |
| Initial / minimum learning rate | 10<sup>−3</sup> / 10<sup>−5</sup> |
| Learning-rate scheduler | ReduceLROnPlateau (factor 0.95, patience 9) |
| Batch size | 1536 |
| Maximum epochs | 500 |
| Input prior | Autoregressive Gaussian (AR1) |
| KL / L2 warmup | 0–80 epochs (linear) |
| Per-area latent dimension *d* | 9 |
| Shared input-space dimension *p<sub>a</sub>* | 64 (M1), 96 (PMd) |
| Flow output dimension *m<sub>a</sub>* | 20 (M1), 40 (PMd) |
| Fisher penalty scale | 10<sup>−4</sup> |
| Fisher SNR target *s*<sup>⋆</sup> | 3 |
| KL scale, controller inputs *κ*<sub>co</sub> | 6 × 10<sup>−5</sup> |
| KL scale, communications *κ*<sub>com</sub> | 0 |
| KL scale, initial conditions *κ*<sub>ic</sub> | 0 |
| MMD scale *κ*<sub>MMD</sub> | 10<sup>−2</sup> |
| MMD sample size | 2048 latent states per session per step |
| MMD kernel widths *cσ* | *c* = 0.5, 1, 2; *σ* = median pairwise distance, recomputed each step |

#### Session-alignment read-in and read-out initialization

Session-alignment read-in matrices were initialized by principal components regression (Pandarinath et al. 2018). For each area, held-in spike counts from eligible subsequent reaches were averaged within the *C* = 8 direction conditions aligned to target onset over *T* time points. These condition averages were Gaussian-smoothed with a standard deviation of 50 ms and concatenated across sessions, yielding a matrix of shape (*H*_1_ + … + *H*_s_) × *CT*, where *H*_s_ is the number of held-in units in session *s*. We performed principal components analysis on this matrix to reduce it to *p*_*a*_ principal components, equal to the dimensionality of the input space, yielding a *p*_*a*_ × *CT* matrix of principal component scores shared by all sessions. For each session, we then linearly regressed these scores against that session’s *H*_*s*_ × *CT* matrix of condition-averaged activity. The regression coefficients and intercepts, which best reconstruct the shared scores from the session’s units, formed its *p*_*a*_ × *H*_*s*_ read-in matrix and its *p*_*a*_-dimensional bias.

Session-alignment read-out matrices were obtained in a similar fashion from the condition-averaged activity of all held-in and held-out units. Firing rates were estimated by smoothing each unit’s spike counts and then averaging within four types of conditions: the direction of the first reach of each trial and the direction of subsequent reaches (each resolved in time), hand position, and hand speed, yielding an (*N*_1_ + … + *N*_*S*_) × *K* matrix, where *N*_*s*_ is the number of units in session *s* and *K* the number of conditions. The condition-averaged rates were passed through the inverse of the rate link, *ρ*^−1^, to obtain pre-activations, and each unit’s mean

pre-activation was subtracted. Before singular value decomposition, each of the four behavioral-condition blocks of the mean-centered pre-activations was scaled to unit Frobenius norm across all sessions. The resulting matrix was reduced by singular value decomposition to *m*_*a*_ components, equal to the output dimensionality of the flow readout, yielding an *m*_*a*_ × *K* shared basis and an (*N*_1_ + … + *N*_*S*_) × *m*_*a*_ matrix of loadings (the left singular vectors scaled by the singular values). For each session *s*, the corresponding *N*_*s*_ × *m*_*a*_ block of loadings, with each row rescaled to unit length, served as its read-out matrix, and the units’ mean pre-activations served as its *N*_*s*_-dimensional bias.

The session-alignment read-in/read-out matrices and biases seed the model with a correspondence across sessions. Because that correspondence is defined by behavioral conditions, the alignment is supervised, although behavior does not enter the training objective.

#### Latent distribution matching

To encourage latent states inferred from different sessions to follow a common distribution, we added a maximum mean discrepancy (MMD) penalty (Gretton et al. 2012) to the training objective that evaluated differences between the distributions of latent activity between every pair of sessions,

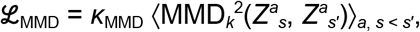

where *κ*_MMD_ = 10^−2^ is the weight of the penalty in the training objective, *Z*^*a*^_*s*_ is a random sample of 2,048 generator states of area *a*, drawn from the windows of session *s* in each minibatch, and ⟨·⟩ denotes the average over areas and pairs of sessions. For two samples *X* and *Y*, the squared MMD is

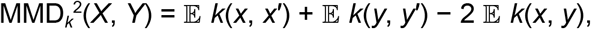

where *x* and *x*′ are states from *X, y* and *y*′ are states from *Y*, and each expectation was estimated by averaging the kernel over all pairs of states in the samples. The kernel *k* measures the similarity of two latent states and was a mixture of three Gaussian kernels,

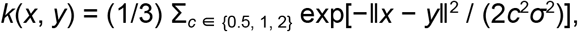

where ‖*x* − *y*‖ is the Euclidean distance between the two states and *cσ* is the width of each kernel. The base width *σ* was set to the median distance between all pairs of states in the combined sample *X* ∪ *Y*, recomputed for every pair of sessions at every training step and treated as a constant when computing gradients. The three kernels therefore compare the two distributions at half, one and two times the typical distance between latent states, so the penalty is sensitive to differences at both fine and coarse scales. Unlike the session-alignment read-in and read-out, this term uses no behavioral information.

#### Training

The multi-session model was trained as described in Training, with the parameters listed in Table 7. Inferred external inputs were regularized toward an AR(1) Gaussian prior with learned per-dimension time constants and innovation variances. Each minibatch contained an equal number of windows from every session, so that every gradient step, and every MMD estimate, involved all sessions. The Fisher-information penalty encouraged local sensitivity of predicted spike counts to latent perturbations and was evaluated on latent states pooled across sessions using the final session’s readout. The checkpoint with the lowest smoothed validation reconstruction loss was used for all analyses.

#### Single-trial and condition-averaged latent trajectories

Latent trajectories (Fig. 6C) of the M1 and PMd generator states and of the PMd→M1 communication signal were computed from the posterior means for every reach, from 0 to 500 ms after target onset, and averaged within each session over the reaches of each of the eight direction conditions. For visualization, a single three-dimensional principal components basis per compartment was fitted to the condition-averaged trajectories of all five sessions together, and single-trial and condition-averaged trajectories were projected onto it, so that all sessions are shown in a common coordinate frame and at a common scale.

#### Cross-session velocity decoding

To test whether the relationship between latent states and behavior was preserved across sessions (Fig. 6D), we fit ridge-regression decoders from each area’s generator state to hand velocity over the first 500 ms after target onset, trained them on one session and evaluated them on every session. Decoders were trained and evaluated with 5-fold cross-validation over reaches, with the ridge penalty selected by internal cross-validation. Entry (*i, j*) in each 5 × 5 matrix shows the *R*^2^ of decoders trained on session *i* and evaluated on session *j’s* out-of-fold reaches.

## Code and data availability

All modeling and analysis code, including the DynISys implementation, preprocessing pipelines, figure-generation scripts, and trained-model checkpoints, will be made publicly available upon publication. The neural data used in this study will also be made publicly available upon publication.

## Acknowledgments

This work was supported by NIH BRAIN/NIDA RF1 DA055667, NIH NINDS/OD DP2 NS127291, and the Air Force Office of Scientific Research award FA95502310727/AR (C.P.), NIH 5F32MH132175 (C.V.), the Simons Foundation through the Simons-Emory International Consortium on Motor Control (C.P., C.V., J.A.P., and J.A.M.), a Banting Postdoctoral Fellowship, a BrainsCAN Postdoctoral Fellowship (CFREF), and a Vector Institute Postgraduate Affiliation (J.A.M.), the Canadian Institutes of Health Research (Project Grant PJT-175010), the Azrieli Foundation through the Collaboration on Motor Planning, Execution and Resilience, and the Canada Research Chair program (J.A.P.).

## Author Contributions

C.V., C.P., and J.A.M. acquired funding for the modeling aspects of this work. J.A.M., J.A.P., and M.K. collected the experimental data. C.V. developed the computational model. C.V. and C.P. designed the analyses, which C.V. conducted with contributions from A.V., T.L., and C.P. C.V., A.V. and T.L. prepared the figures. C.V. wrote the manuscript, with contributions from C.P., A.V., and T.L. J.A.M., J.A.P., and M.K. provided feedback on the analyses and manuscript.

---

**Extended Data Figure 1.**
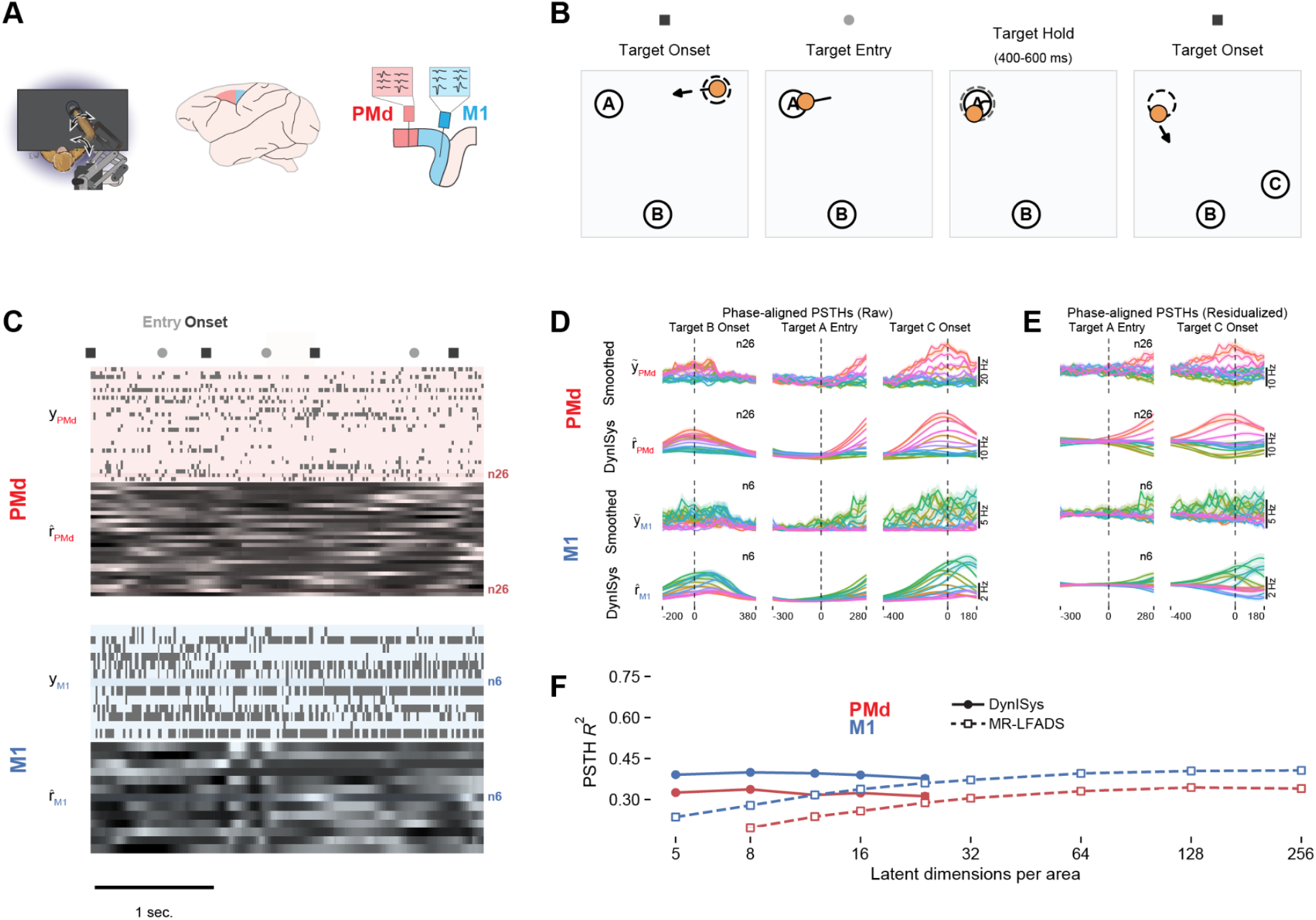
DynISys captures PMd/M1 population activity patterns during a sequential reaching task (Monkey M). **(A)** Kinarm behavioral apparatus and PMd/M1 recording-array locations. **(B)** Target (unfilled circles) and hand position (orange circle) across task phases of a single trial: target onset (target B appears, reach to target A begins), target entry (hand enters A), target hold, and second target onset (target C appears, reach to B begins). **(C)** Example PMd/M1 spike rasters (top) and matching DynISys-predicted firing-rate heatmaps (bottom) for the same stitched trial window, showing single-trial, per-channel rate agreement with observed spiking. **(D)** PMd/M1 PSTHs aligned to target B onset, target A entry, and target C onset, comparing observed and DynISys-predicted firing rates across phases. Reach angle indicated by colorwheel. **(E)** Same as (D), with PSTHs computed after removing components of the neural activity that were predictive of the current target location using linear models. **(F)** PSTH R^2^ for DynISys and MR-LFADS as a function of latent dimensionality.

**Extended Data Figure 2.**
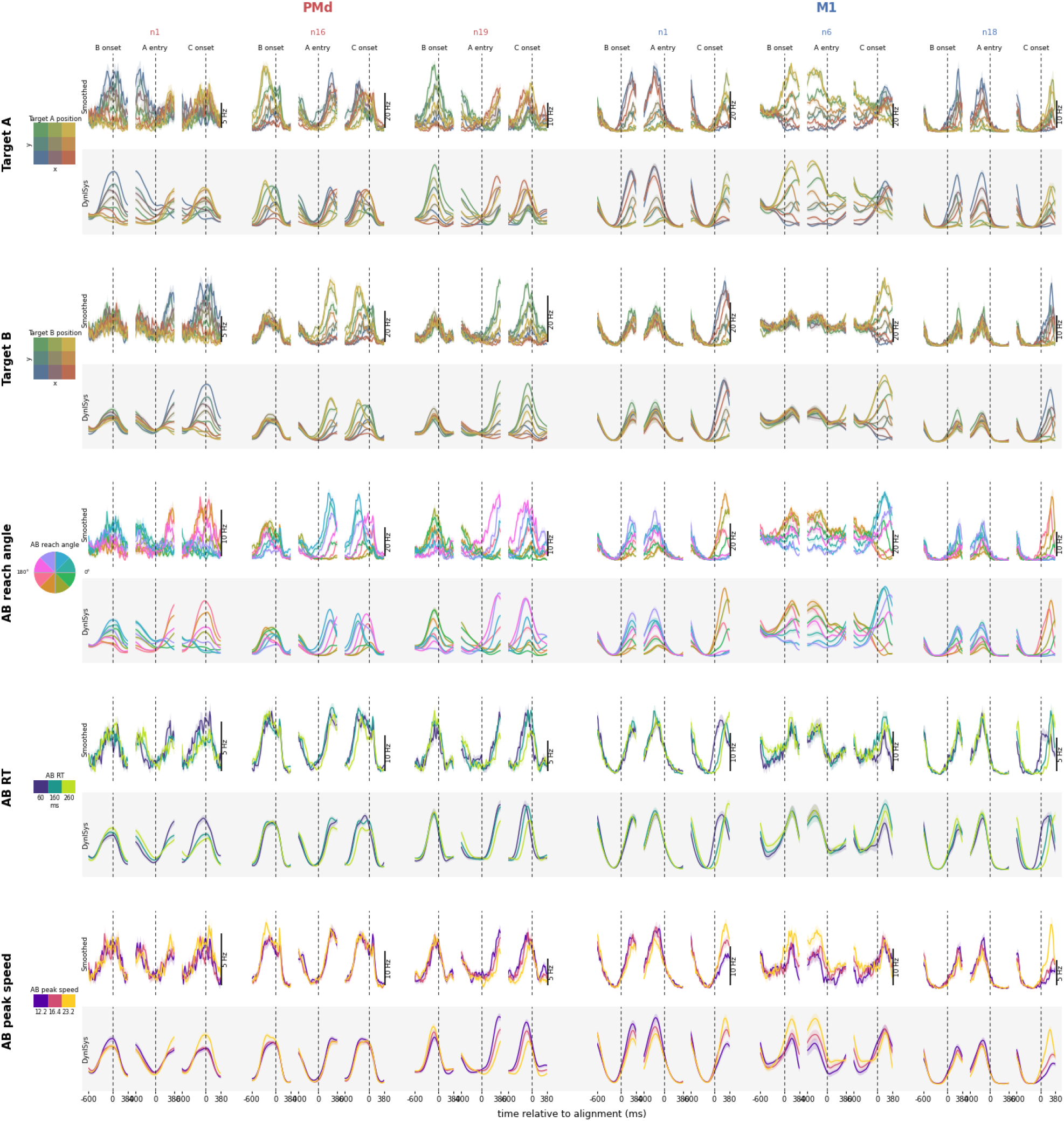
DynISys predicted firing rates capture single neuron diversity and complex encoding of multiple behavioral features (Monkey P). Causally smoothed spiking activity (upper rows) and DynISys-inferred firing rate (lower rows, grey) PSTHs for example PMd and M1 neurons. Columns are grouped by neuron. Left: PMd neurons. Right: M1 neurons. Three task-epoch alignments per neuron: B onset, A entry, and C onset. The time axis is relative to each alignment event (ms). Rows depict PSTHs, with trials grouped by a different feature. *Row 1:* Target A position. *Row 2:* Target B position, colored by a two-dimensional (x, y) position key (3×3 spatial grid, bilinear colormap); *Row 3:* A→B reach angle, split into 8 circular bins indicated by colorwheel matching Fig. 2. *Row 4*: A→B reaction time, and *Row 5*: A→B peak hand speed. Reaction time and peak hand speed groups were generated by splitting trials into tertiles by the respective feature and colored by a distinct sequential colormap. Each group’s median value is indicated beneath its colorbar. Traces denote within-group mean firing rate for each time bin (shaded band, s.e.m.). Vertical scale bars (right of each block) denote firing rate; y-axis is shared across the three alignments within each neuron/variable block but varies between rows and neurons.

**Extended Data Figure 3.**
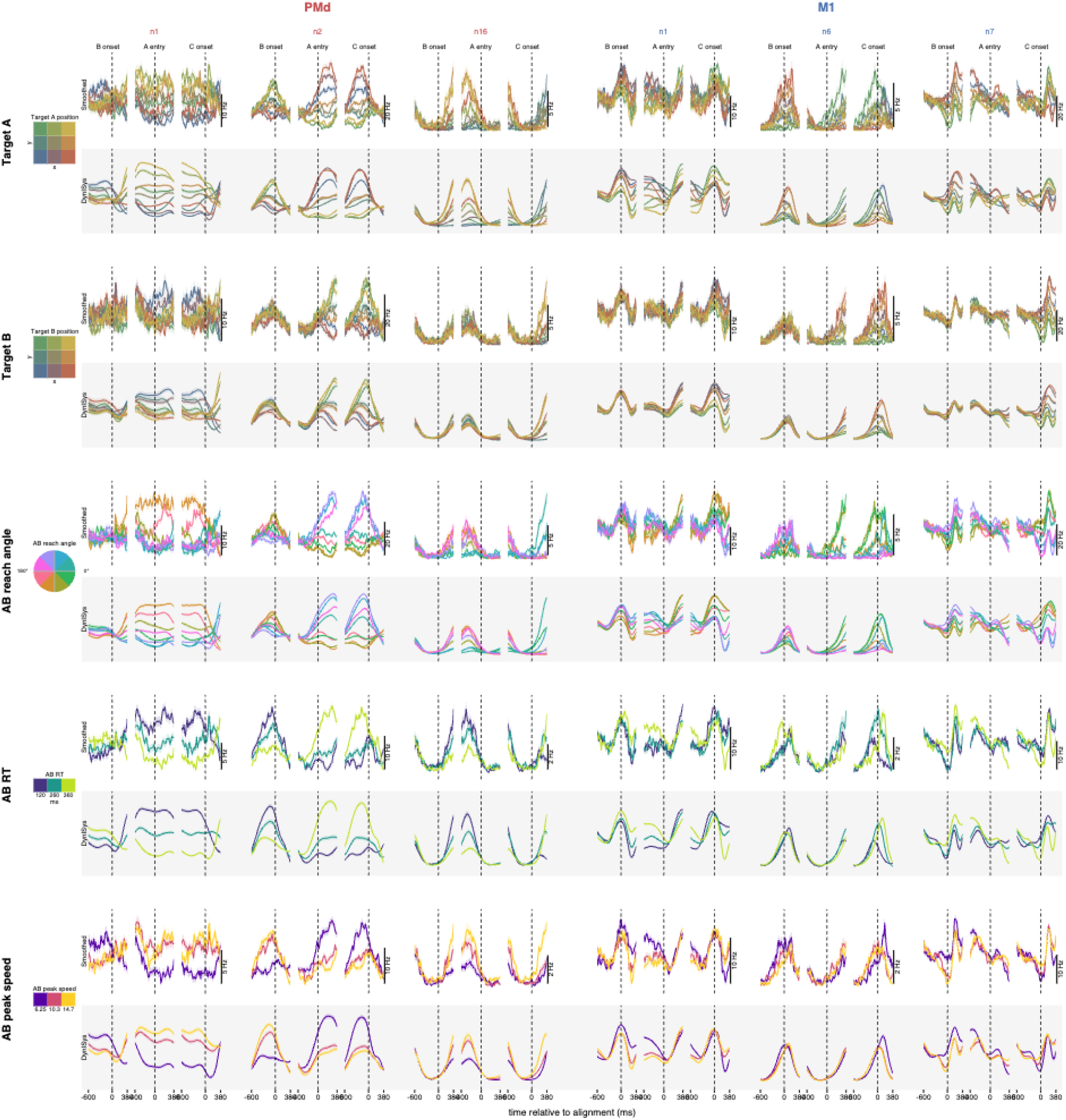
DynISys-predicted firing rates capture single neuron diversity and complex encoding of multiple behavioral features (Monkey M). As in Extended Data Figure 2, but for Monkey M.

**Extended Data Figure 4.**
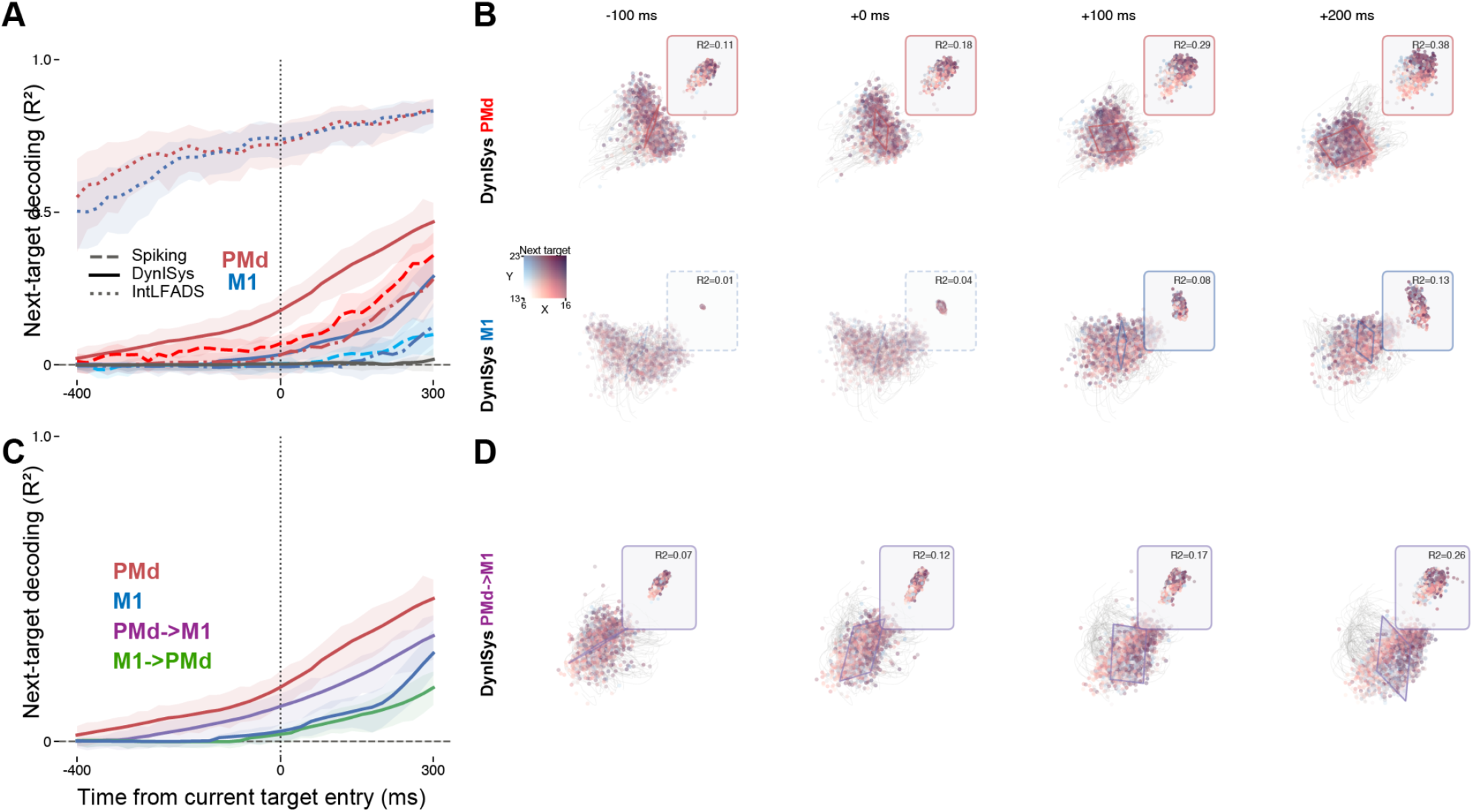
DynISys accurately infers the time course of target encoding within each area and transmission of target information across areas (Monkey M). **(A)** Time-resolved next-target decoding from PMd latents, PMd smoothed spikes, M1 latents, M1 smoothed spikes, and IntLFADS controls, aligned to current-target entry. Shaded areas indicate bootstrapped 95% confidence intervals for mean decoding performance. **(B)** State-space snapshots of PMd and M1 DynISys latents (top-3 PCs) at four times around current-target entry, each with a fitted linear decoding plane for next-target position (colored by target, R^2^ noted); inset shows decoded-vs-true target projection per time slice. **(C)** Time-resolved next-target decoding from PMd→M1 communication vs. the M1→PMd control direction (plus PMd/M1 latents for reference). **(D)** Same state-space/plane treatment as (B) for the PMd→M1 communication channel.

**Extended Data Table 1.** Architectural differences between DynISys, IntLFADS, and MR-LFADS.

| **Feature/Model** | **DynISys** | **IntLFADS** | **MR-LFADS** |
|---|---|---|---|
| **Generator** | Neural ODE | RNN | RNN |
| **Communication** | Latent | Latent | Neural |
| **Readout** | Injective nonlinear | Linear | Linear |
| **Initial-condition inf.** | Causal | Acausal | Causal |

**Extended Data Figure 5.**
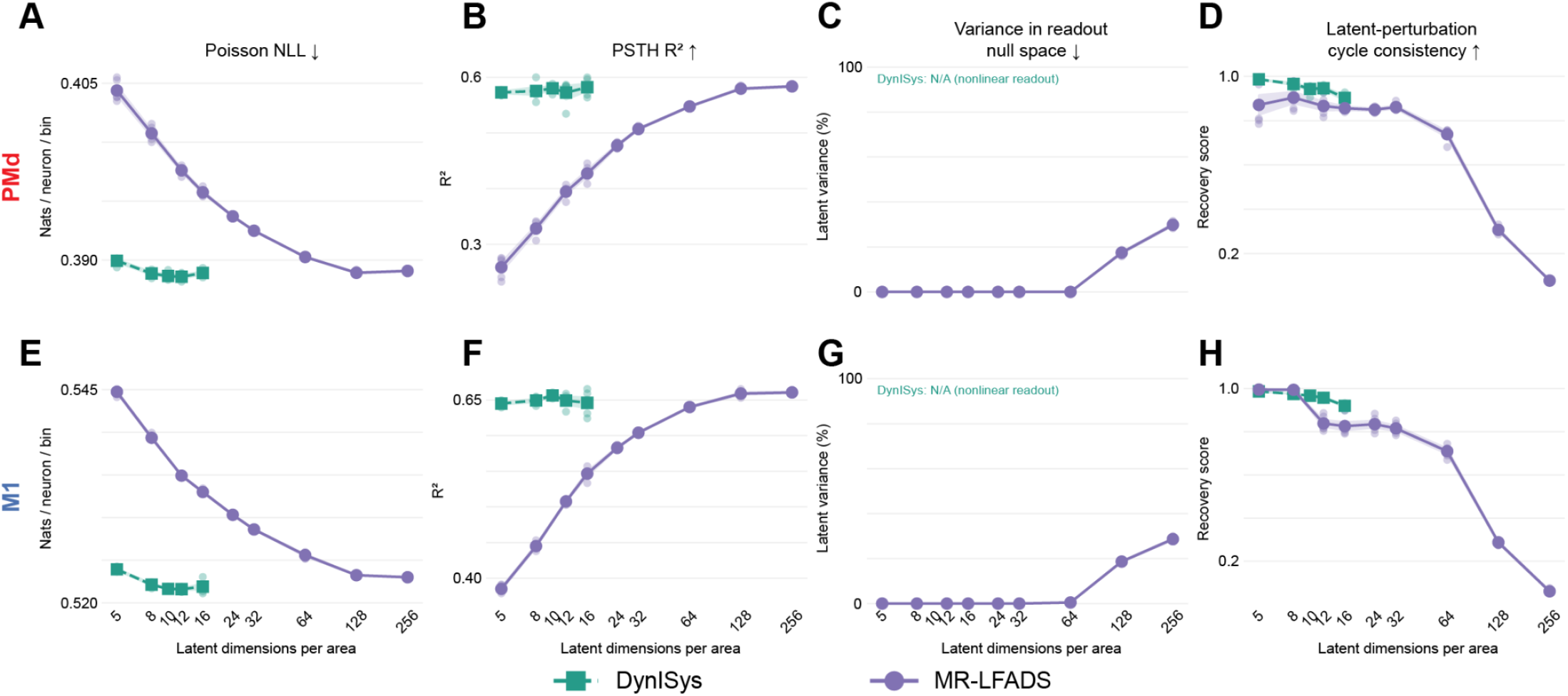
Accurate neural reconstruction can coexist with poor recovery of latent perturbations in MR-LFADS. **(A–D)** Comparison of MR-LFADS (purple) and DynISys (teal) across latent dimensionalities for PMd: **(A)** validation Poisson negative log-likelihood, **(B)** PSTH R^2^ computed from training and validation trials, (**C)** percentage of validation latent variance in the null space of the MR-LFADS linear readout, and **(D)** latent-perturbation cycle-consistency score, measuring recovery of independent latent displacements after passing perturbed states through the readout and a learned inverse. **(E–H)** Same as **(A–D)**, for M1. At larger latent dimensionalities, MR-LFADS maintains accurate neural reconstruction while null-space variance increases and latent-perturbation recovery declines. The linear null-space diagnostic is not applicable to the nonlinear DynISys readout. Lines show means across five training seeds, shaded bands indicate ±SEM, and points show individual seeds.

**Extended Data Figure 6.**
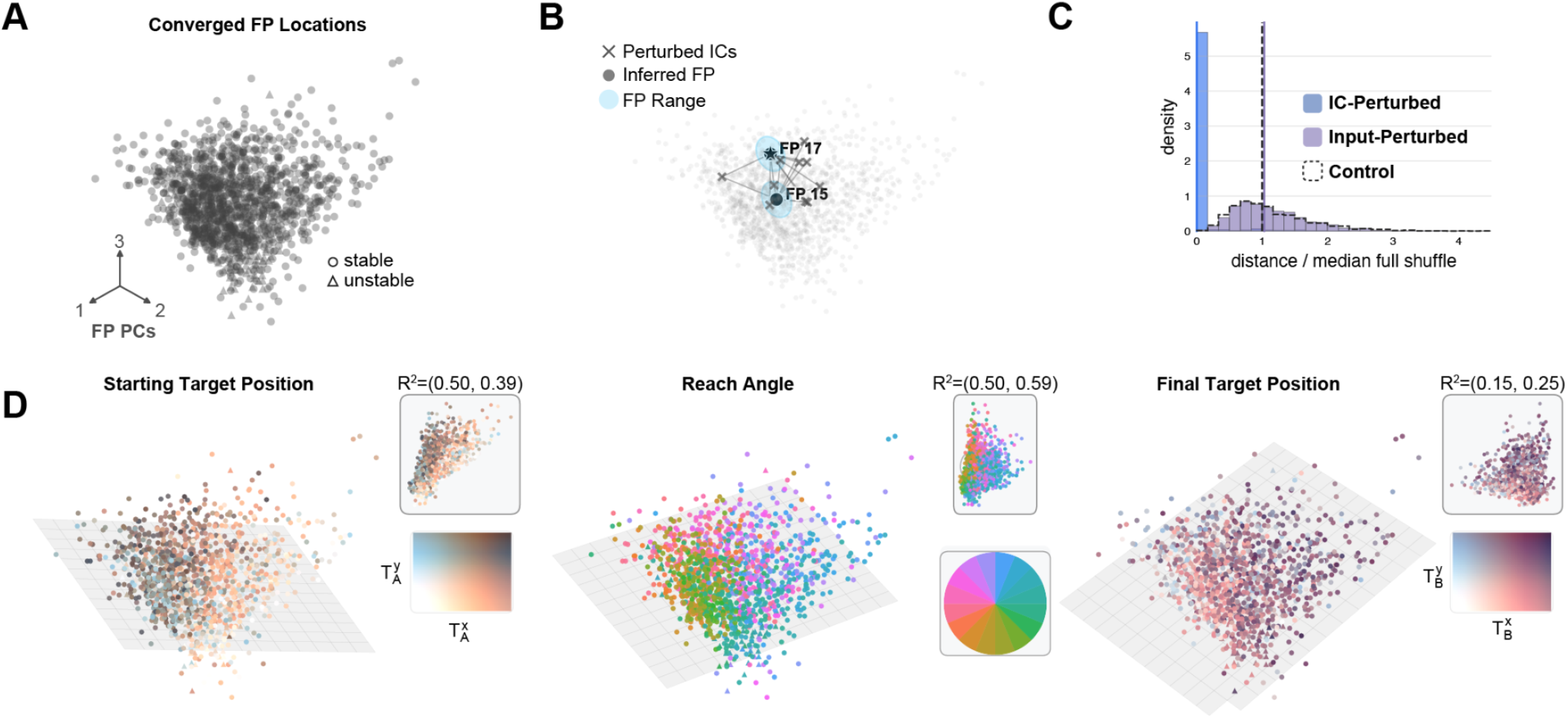
M1 dynamics exhibit input-modulated fixed points that reliably encode features of the upcoming reach (Monkey M). **(A)** Top 3 PCs of inferred M1 fixed points from optimization with each trial’s own inputs and initial condition. Fixed points converged below cutoff (q-value < 1e-6) on 1214/1313 trials. Of converged fixed points, 1198/1214 (>98%) were stable, indicated with circles; unstable fixed points indicated by triangles. **(B)** Example fixed point descents, computed using two trials’ inferred inputs and 10 randomly selected initial conditions. Converged fixed point locations are highlighted with cyan circles. Example fixed point optimizations tend to converge to locations organized by the external input, rather than initial conditions. **(C)** Histogram of latent distances from same-trial fixed point locations (panel A) to fixed points computed with the same trial’s input (cyan) or initial condition (purple). Shuffled control (distance from original fixed point to fixed point inferred by random trial’s IC and input) shown as dashed line. Static input fixed points tend to be much closer to the same-trial fixed point than static IC fixed points. **(D)** Inferred fixed point locations encode features of upcoming reach. Each column shows same-trial fixed point locations color coded by features of upcoming reach. Colormap of feature and decoder output / performance shown as insets in each column (bottom, top insets, respectively). Left: Starting target position. Middle: Reach angle. Right: Final target position.

**Extended Data Figure 7.**
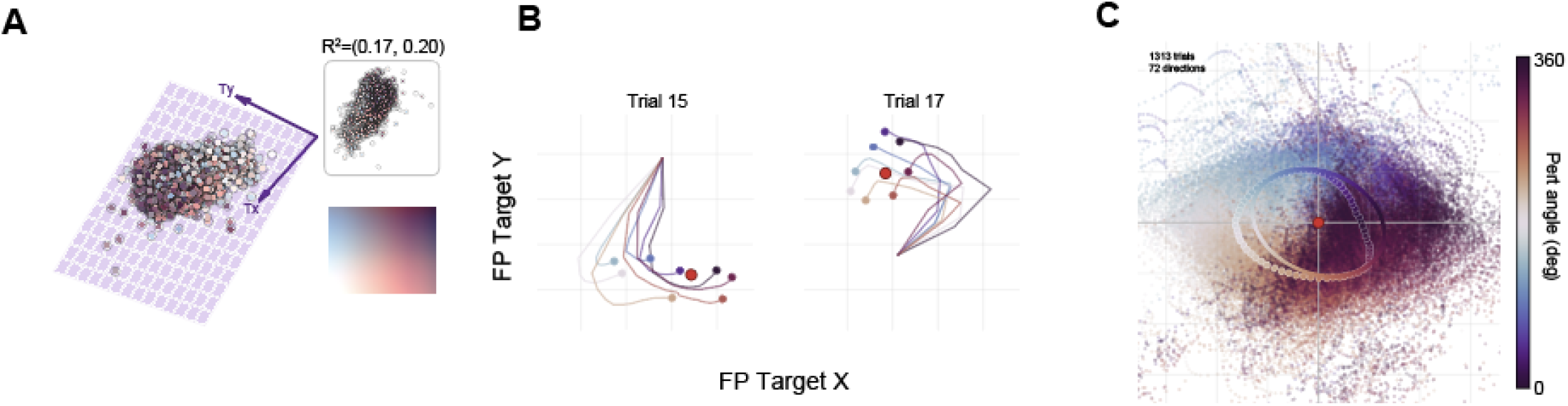
Simulated PMd perturbations identify functional role of inferred PMd inputs to M1 (Monkey M). **(A)** PMd→M1 communication channel activity ĉ_*PMd*−>*M*1_ projected onto its top 3 PCs, colored by upcoming reach target (X/Y, see inset). A linear decoder (purple plane) maps this channel onto the next target; inset shows decoded vs. true target performance R^2^. **(B)** For two example trials, converged M1 fixed points obtained by descending the flow field under systematically perturbed PMd→M1 inputs (colored by perturbation direction), decoded into target space. Red marker = unperturbed fixed point; yellow marker = FP finding initial condition. **(C)** Across all trials, fixed point displacement (perturbed − unperturbed, decoded target space) as function of perturbation direction (color, 0-360°), with per-direction means and best-fit response ellipse. M1 fixed points shift in the direction predicted by the communication channel’s decoded target.

**Extended Data Figure 8.**
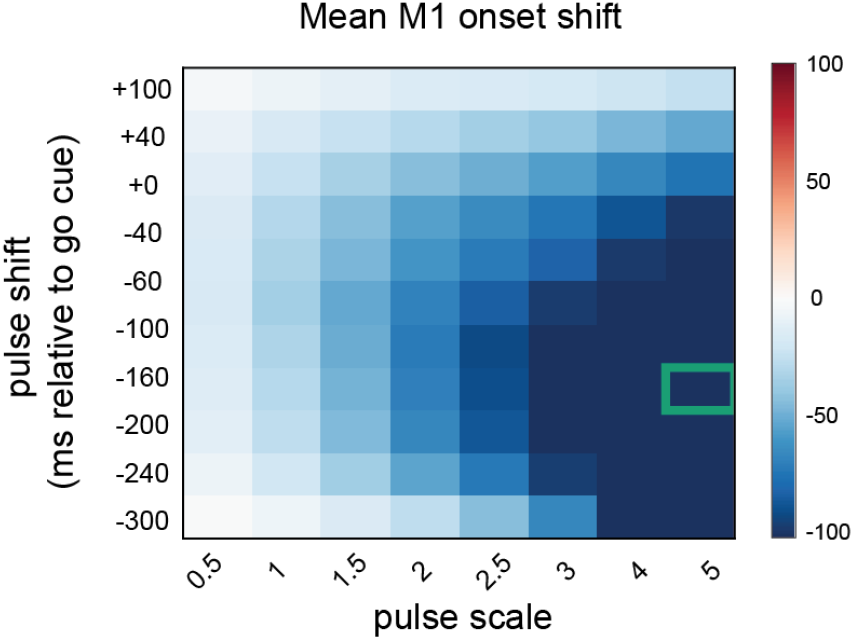
Pulse parameter dependence of M1 onset shift induced by input perturbation. Mean shift in M1 movement onset time (color scale, ms) as a function of perturbation pulse scale (x-axis) and pulse timing relative to the go cue (y-axis). Onset shifts become larger (more negative) with increasing pulse scale. Green box indicates the parameter combination used for the perturbation shown in the main text (Fig. 5E-G).

**Extended Data Figure 9.**
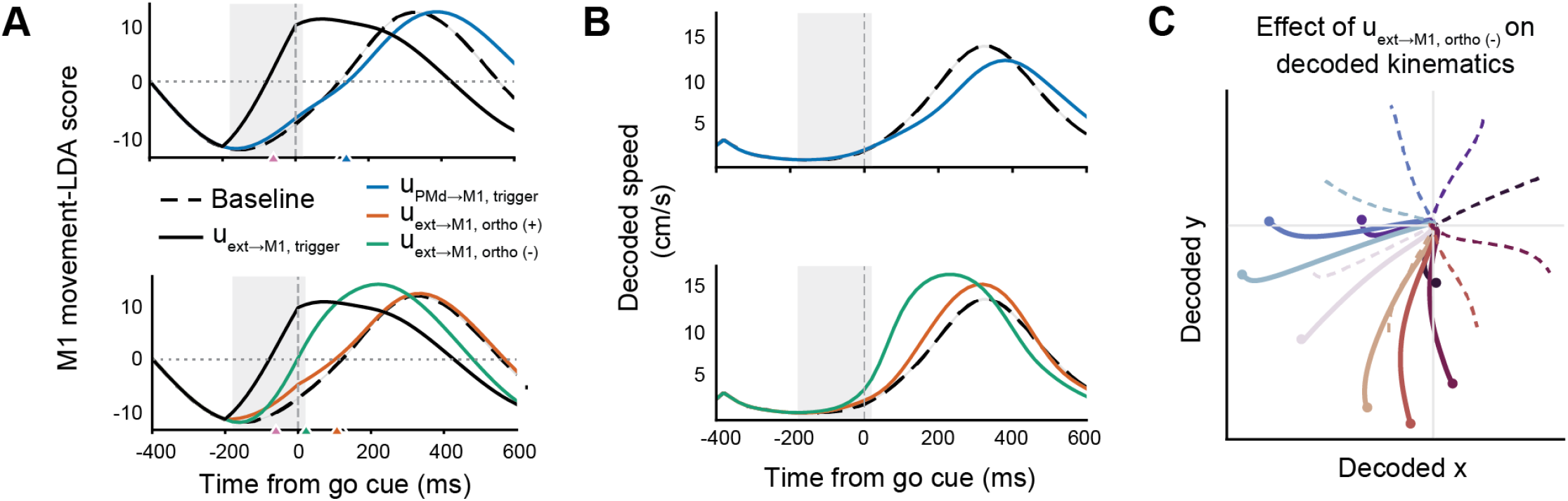
Input perturbations reveal distinct effects on movement timing and direction. **(A)** M1 movement-LDA scores, aligned to the go cue, for baseline trials (dashed black) versus perturbed trials. *Top:* trigger-axis perturbation (solid black) advances M1 neural onset by 182.2 ± 2.3 ms, whereas perturbing along the condition-invariant component of PMd→M1 (blue) input delays onset by 17.1 ± 2.2 ms. *Bottom:* perturbing the external input along the highest variance direction orthogonal to the trigger axis identified in Fig. 5E. Positive (orange) and negative (green) pulses along this axis shift M1 neural onset by −15.6 ± 2.9 ms and −100.0 ± 2.1 ms, respectively, relative to baseline (mean ± SEM; negative shifts indicate earlier onset). Triangles indicate mean per-trial neural onset times. **(B)** Corresponding decoded speed profiles (cm/s) for baseline and control perturbations. Grey shading in A and B marks the pulse window (−180 to +20 ms relative to the go cue; pulse scale 5). **(C)** Decoded trajectories following orthogonal (−) stimulation (colored by target direction), overlaid on baseline trajectories (dashed). Orthogonal (−) stimulation advances neural onset while also altering decoded movement direction. Dots mark trajectory endpoints at +600 ms.

## References

1. Chen, Ricky T. Q., Yulia Rubanova, Jesse Bettencourt, and David Duvenaud. 2018. “Neural Ordinary Differential Equations.” Advances in Neural Information Processing Systems (NeurIPS).

2. Churchland, Mark M., John P. Cunningham, Matthew T. Kaufman, et al. 2012. “Neural Population Dynamics During Reaching.” Nature 487 (7405): 51–56. 10.1038/nature11129.

3. Churchland, M. M., J. P. Cunningham, M. T. Kaufman, S. I. Ryu, and K. V. Shenoy. 2010. “Cortical Preparatory Activity: Representation of Movement or First Cog in a Dynamical Machine?” Neuron 68 (3): 387–400. 10.1016/j.neuron.2010.09.015.

4. Driscoll, Laura N., Krishna Shenoy, and David Sussillo. 2024. “Flexible Multitask Computation in Recurrent Networks Utilizes Shared Dynamical Motifs.” Nature Neuroscience 27 (7): 1349–1363.

5. Durstewitz, Daniel, Georgia Koppe, and Max Ingo Thurm. 2023. “Reconstructing Computational System Dynamics from Neural Data with Recurrent Neural Networks.” Nature Reviews Neuroscience 24 (11): 693–710.

6. Gao, Yuanjun, Evan W. Archer, Liam Paninski, and John P. Cunningham. 2016. “Linear Dynamical Neural Population Models through Nonlinear Embeddings.” Advances in Neural Information Processing Systems 29.

7. Glaser, Joshua I., Matthew G. Perich, Pavan Ramkumar, Lee E. Miller, and Konrad P. Kording. 2018. “Population Coding of Conditional Probability Distributions in Dorsal Premotor Cortex.” Nature Communications 9: 1788. 10.1038/s41467-018-04062-6.

8. Glaser, Joshua I., Matthew R. Whiteway, John P. Cunningham, Liam Paninski, and Scott W. Linderman. 2020. “Recurrent Switching Dynamical Systems Models for Multiple Interacting Neural Populations.” Advances in Neural Information Processing Systems (NeurIPS).

9. Gokcen, Evren, Anna I. Jasper, João D. Semedo, et al. 2022. “Disentangling the Flow of Signals Between Populations of Neurons.” Nature Computational Science 2 (8): 512–25. 10.1038/s43588-022-00282-5.

10. Golub, Matthew D., and David Sussillo. 2018. “FixedPointFinder: A Tensorflow Toolbox for Identifying and Characterizing Fixed Points in Recurrent Neural Networks.” Journal of Open Source Software 3 (31): 1003.

11. Gretton, Arthur, Karsten M. Borgwardt, Malte J. Rasch, Bernhard Schölkopf, and Alexander Smola. 2012. “A Kernel Two-Sample Test.” Journal of Machine Learning Research 13: 723–773.

12. Inagaki, Hidehiko K., Lorenzo Fontolan, Sandro Romani, and Karel Svoboda. 2019. “Discrete Attractor Dynamics Underlies Persistent Activity in the Frontal Cortex.” Nature 566 (7743): 212–17. 10.1038/s41586-019-0919-7.

13. Inagaki, Hidehiko K., Susu Chen, Margreet C. Ridder, et al. 2022. “A Midbrain-Thalamus-Cortex Circuit Reorganizes Cortical Dynamics to Initiate Movement.” Cell 185 (6): 1065–1081.e23.

14. Karniol-Tambour, Orren, David M. Zoltowski, E. Mika Diamanti, et al. 2024. “Modeling State-Dependent Communication between Brain Regions with Switching Nonlinear Dynamical Systems.” International Conference on Learning Representations (ICLR), 27010–27030.

15. Kaufman, Matthew T., Mark M. Churchland, Stephen I. Ryu, and Krishna V. Shenoy. 2014. “Cortical Activity in the Null Space: Permitting Preparation without Movement.” Nature Neuroscience 17 (3): 440–448.

16. Kaufman, Matthew T., Jeffrey S. Seely, David Sussillo, et al. 2016. “The Largest Response Component in the Motor Cortex Reflects Movement Timing but Not Movement Type.” eNeuro 3 (4): ENEURO.0085-16.2016.

17. Kao, J. C. 2019. “Considerations in Using Recurrent Neural Networks to Probe Neural Dynamics.” Journal of Neurophysiology 122 (6): 2504–2521. 10.1152/jn.00467.2018.

18. Kashefi, Mehrdad, Jonathan A. Michaels, Rhonda Kersten, J. C. Lau, Jörn Diedrichsen, and J. Andrew Pruszynski. 2026. “Compositional Neural Dynamics during Reaching.” bioRxiv. Revised January 29, 2026. 10.1101/2025.09.04.674069.

19. Keshtkaran, Mohammad Reza, Andrew R. Sedler, Raeed H. Chowdhury, et al. 2022. “A Large-Scale Neural Network Training Framework for Generalized Estimation of Single-Trial Population Dynamics.” Nature Methods 19 (12): 1572–77. 10.1038/s41592-022-01675-0.

20. Kim, Timothy D., Thomas Z. Luo, Jonathan W. Pillow, and Carlos D. Brody. 2021. “Inferring Latent Dynamics Underlying Neural Population Activity via Neural Differential Equations.” In Proceedings of the 38th International Conference on Machine Learning, 139:5551–5561. Proceedings of Machine Learning Research. PMLR. https://proceedings.mlr.press/v139/kim21h.html.

21. Kohn, Adam, Ruben Coen-Cagli, Ingmar Kanitscheider, and Alexandre Pouget. 2016. “Correlations and Neuronal Population Information.” Annual Review of Neuroscience 39: 237–256.

22. Linderman, Scott W., Andrew C. Miller, Ryan P. Adams, et al. 2016. “Recurrent Switching Linear Dynamical Systems.” 1610.08466.

23. Liu, Belle, Jacob Sacks, and Matthew D. Golub. 2025. “Accurate Identification of Communication Between Multiple Interacting Neural Populations.” In Proceedings of the 42nd International Conference on Machine Learning, 267:39381–39404. Proceedings of Machine Learning Research. PMLR. https://proceedings.mlr.press/v267/liu25bh.html.

24. Macke, Jakob H., Lars Buesing, John P. Cunningham, et al. 2011. “Empirical Models of Spiking in Neural Populations.” Advances in Neural Information Processing Systems 24.

25. Maheswaranathan, Niru, Alex H. Williams, Matthew D. Golub, et al. 2019. “Universality and Individuality in Neural Dynamics across Large Populations of Recurrent Networks.” Advances in Neural Information Processing Systems 32.

26. Marraffa, A., R. Krause, V. Mante, et al. 2026. “Data-Driven Reduced Modeling of Neural Dynamics.” Nature Communications. 10.1038/s41467-026-75924-7.

27. Mastrogiuseppe, Francesca, and Srdjan Ostojic. 2018. “Linking Connectivity, Dynamics, and Computations in Low-Rank Recurrent Neural Networks.” Neuron 99 (3): 609–623.e29.

28. Michaels, J. A., M. Kashefi, J. Zheng, et al. 2025. “Sensory Expectations Shape Neural Population Dynamics in Motor Circuits.” Nature 648 (8094): 668–77. 10.1038/s41586-025-09690-9.

29. Oby, Emily R., Alan D. Degenhart, Erinn M. Grigsby, et al. 2025. “Dynamical Constraints on Neural Population Activity.” Nature Neuroscience 28 (2): 383–393.

30. Pandarinath, Chethan, Daniel J. O’Shea, Jasmine Collins, et al. 2018. “Inferring Single-Trial Neural Population Dynamics Using Sequential Auto-Encoders.” Nature Methods 15 (10): 805–15. 10.1038/s41592-018-0109-9.

31. Perich, Matthew G., Charlotte Arlt, Sofia Soares, et al. 2026. “Inferring Brain-Wide Interactions Using Data-Constrained Recurrent Neural Network Models.” Neuron. 10.1016/j.neuron.2026.07.016.

32. Perich, Matthew G., Lee E. Miller, Mehdi Azabou, and Eva L. Dyer. 2025. Long-Term Recordings of Motor and Premotor Cortical Spiking Activity During Reaching in Monkeys. Version 0.250122.1735. DANDI Archive. 10.48324/dandi.000688/0.250122.1735.

33. Rezende, Danilo Jimenez, and Shakir Mohamed. 2015. “Variational Inference with Normalizing Flows.” International Conference on Machine Learning (ICML), 1530–38.

34. Sauerbrei, Britton A., Jian-Zhong Guo, Jeremy D. Cohen, et al. 2020. “Cortical Pattern Generation during Dexterous Movement Is Input-Driven.” Nature 577 (7790): 386–391.

35. Schimel, Marine, Ta-Chu Kao, Kristopher T. Jensen, and Guillaume Hennequin. 2022. “iLQR-VAE: Control-Based Learning of Input-Driven Dynamics with Applications to Neural Data.” International Conference on Learning Representations (ICLR).

36. Scott, Stephen H. 1999. “Apparatus for Measuring and Perturbing Shoulder and Elbow Joint Positions and Torques during Reaching.” Journal of Neuroscience Methods 89 (2): 119–127.

37. Sedler, Andrew R., Christopher Versteeg, and Chethan Pandarinath. 2023. “Expressive Architectures Enhance Interpretability of Dynamics-Based Neural Population Models.” Neurons, Behavior, Data Analysis, and Theory, March, 1–22. 10.51628/001c.73987.

38. Semedo, João D., Amin Zandvakili, Christian K. Machens, Byron M. Yu, and Adam Kohn. 2019. “Cortical Areas Interact Through a Communication Subspace.” Neuron 102 (1): 249–259.e4. 10.1016/j.neuron.2019.01.026.

39. Shah, Nishal P., Benyamin Abramovich Krasa, Erin Kunz, et al. 2025. “Improved Interpretability in LFADS Models Using a Learned, Context-Dependent Per-Trial Bias.” bioRxiv.

40. Shenoy, Krishna V., Maneesh Sahani, and Mark M. Churchland. 2013. “Cortical Control of Arm Movements: A Dynamical Systems Perspective.” Annual Review of Neuroscience 36: 337–59. 10.1146/annurev-neuro-062111-150509.

41. Sussillo, David, and Omri Barak. 2013. “Opening the Black Box: Low-Dimensional Dynamics in High-Dimensional Recurrent Neural Networks.” Neural Computation 25 (3): 626–49. 10.1162/NECO_a_00409.

42. Vahidi, Parsa, Omid G. Sani, and Maryam M. Shanechi. 2025. “BRAID: Input-Driven Nonlinear Dynamical Modeling of Neural-Behavioral Data.” International Conference on Learning Representations (ICLR), 2940–2969.

43. Versteeg, C., J. D. McCart, M. Ostrow, D. M. Zoltowski, C. B. Washington, L. Driscoll, et al. 2026. “Computation-through-Dynamics Toolkit: Simulated Datasets and Quality Metrics for Dynamical Models of Neural Activity.” PLOS Computational Biology 22 (10): e1014755. 10.1371/journal.pcbi.1014755.

44. Versteeg, Christopher, Andrew R. Sedler, Jonathan D. McCart, and Chethan Pandarinath. 2024. “Expressive Dynamics Models with Nonlinear Injective Readouts Enable Reliable Recovery of Latent Features from Neural Activity.” In Proceedings of the 2nd NeurIPS Workshop on Symmetry and Geometry in Neural Representations, 228:255–278. Proceedings of Machine Learning Research. PMLR. https://proceedings.mlr.press/v228/versteeg24a.html.

45. Vyas, Saurabh, Matthew D. Golub, David Sussillo, and Krishna V. Shenoy. 2020. “Computation Through Neural Population Dynamics.” Annual Review of Neuroscience 43: 249–75. 10.1146/annurev-neuro-092619-094115.

46. Wang, E. Y., P. G. Fahey, Z. Ding, et al. 2025. “Foundation Model of Neural Activity Predicts Response to New Stimulus Types.” Nature 640: 470–477. 10.1038/s41586-025-08829-y.

47. Zhu, Feng, Harrison A. Grier, Raghav Tandon, et al. 2022. “A Deep Learning Framework for Inference of Single-Trial Neural Population Dynamics from Calcium Imaging with Subframe Temporal Resolution.” Nature Neuroscience 25 (12): 1724–1734.

48. Zimnik, Andrew J., and Mark M. Churchland. 2021. “Independent Generation of Sequence Elements by Motor Cortex.” Nature Neuroscience 24 (3): 412–24. 10.1038/s41593-021-00798-5.

